# Commensal myeloid crosstalk in neonatal skin regulates cutaneous type 17 inflammation

**DOI:** 10.1101/2023.09.29.560039

**Authors:** Miqdad O. Dhariwala, Ricardo O. Carale, Andrea M. DeRogatis, Henry Rodriguez-Valbuena, Joy N. Okoro, Christina Ekstrand, Antonin Weckel, Victoria M. Tran, Irek Habrylo, Rio Barrere-Cain, Oluwasunmisola T. Ojewumi, Allison E. Tammen, Jessica Tsui, Saba Shaikh, Shivaram Yellamilli, Ahmed K. Aladhami, John M. Leech, Geil R. Merana, Kamir J. Hiam-Galvez, Joanna Halkias, James Gardner, Rachel Rutishauser, Gabriela K. Fragiadakis, Matthew H. Spitzer, Alexis J. Combes, Tiffany C. Scharschmidt

## Abstract

Early life microbe-immune interactions at barrier surfaces have lasting impacts on the trajectory towards health versus disease. Monocytes, macrophages and dendritic cells are primary sentinels in barrier tissues, yet the salient contributions of commensal-myeloid crosstalk during tissue development remain poorly understood. Here, we identify that commensal microbes facilitate accumulation of a population of monocytes in neonatal skin. Transient postnatal depletion of these monocytes resulted in heightened IL-17A production by skin T cells, which was particularly sustained among CD4^+^ T cells and sufficient to exacerbate inflammatory skin pathologies. Neonatal skin monocytes were enriched in expression of negative regulators of the IL-1 pathway. Functional in vivo experiments confirmed a key role for excessive IL-1R1 signaling in T cells as contributing to the dysregulated type 17 response in neonatal monocyte-depleted mice. Thus, a commensal-driven wave of monocytes into neonatal skin critically facilitates immune homeostasis in this prominent barrier tissue.

## Introduction

Symbiotic relationships established between commensal microbes and the immune system help lay the foundation for tissue homeostasis at body barrier sites. Such interactions are especially pivotal in the early life window for defining the longer-term trajectory of health versus disease^1,2,3^. Neonatal exposure to skin commensal bacteria has been shown to critically shape the composition and function of cutaneous lymphocytes^4^. Work by our group and others have demonstrated the importance of early microbial cues in recruiting CD4^+^ regulatory T cells (Tregs)^5^ and Mucosal Associated Invariant T (MAIT) lymphocytes to the neonatal skin^6^, cell types which in turn promote enduring commensal-specific immune tolerance and improved wound repair. Comparatively little is known, however, about early life crosstalk between commensal microbes and skin myeloid cells, despite the sentinel role these cells play in sensing microbial cues.

The developing immune system is enriched, by design, in immune regulatory mechanisms to help limit inflammation in response to an onslaught of new antigenic exposures^7,8^. This extends to barrier tissues, where antigen-presenting cells (APCs) have been specifically implicated in promoting tolerance via Treg generation in the perinatal window. For example, arginase-2 producing dendritic cells (DC) in human prenatal tissues^9^, RORγt-expressing antigen-presenting cells in neonatal mesenteric lymph nodes^10^, and CD301b^+^ type 2 DCs in neonatal skin^11^ have all been shown to facilitate Treg generation, the latter two specifically in response commensal antigens. What additional mechanisms and other myeloid cell types might contribute to the regulatory environment of developing barrier tissues remain undefined.

Monocytes are circulating mononuclear myeloid cells constitutively produced in the bone marrow that, upon entering non-lymphoid tissues such as skin, can further differentiate into macrophages and subsets of DCs. Monocytes help initiate and tune immune response via multiple mechanisms including phagocytosis, antigen presentation and cytokine secretion^12^. In mice, functionally distinct monocyte subsets are distinguishable via their respective expression of the chemokine receptors CCR2 and CX3CR1. CCR2^hi^ classical monocytes, are generally recognized for their pro-inflammatory contributions towards initiation of immune response following infection or injury, whereas CX3CR1^hi^ non-classical or patrolling monocytes have described contributions in immune surveillance, tissue-repair and anti-tumor immune responses^13–16^. However, many unanswered questions remain about the composition and function of monocytes in developing barrier tissues, including whether they contribute to the immune regulatory environment and if they differ from their adult counterparts or to those in inflamed tissues.

Here, we set out to better define the early life skin myeloid compartment and its relationship to microbial colonization, uncovering that CCR2^hi^Ly6C^hi^ monocytes are enriched in murine neonatal skin and that commensal microbes play a vital role in their accumulation. We discovered that transient neonatal depletion of these monocytes led to a persistent, dysregulated type 17 inflammatory response among skin T cells. Heightened expression of IL-1 pathway repressors, IL-1Ra and IL-1R2, by neonatal skin monocytes and their maturing progeny as well as restoration of a homeostatic type 17 response in monocyte-depleted mice via neonatal blockade of IL-1 signaling, identified IL-1 modulation as a key axis for monocyte-mediated immune regulation in neonatal skin. Our study reports a novel regulatory immune role for monocytes in neonatal tissues, by which they help establish a trajectory for tissue immune homeostasis.

## Results

### Classical monocytes rapidly accumulate in neonatal skin

To gain a global understanding of the myeloid cell landscape in murine skin during early postnatal development we used Mass Cytometry (CyTOF) to enumerate skin immune cells in wild type (wt) C57BL6 mice during the first month of life (Fig. 1A). We developed a 34 antibody panel heavily focused on myeloid cell markers along with sufficient lymphocyte epitopes to enable basic identification of T cell subsets. Application of this panel to 6-, 15-and 30-day-old back skin revealed stark age-dependent differences as demonstrated by principal component analysis on the CD45^+^ compartment (Fig. 1B, Fig. S1A). Pooled analysis of skin from all three time points identified 26 distinct clusters of immune cells (Fig. 1C, Fig. S1B). Six were identified as lymphocyte subtypes and fifteen were of clear myeloid origin. The remaining five could not be definitively identified based on our antibody panel, but likely included innate lymphoid cells (ILCs)^17^. Multiple clusters were specifically enriched at distinct time points (Fig. 1D-E).

**Figure 1:**
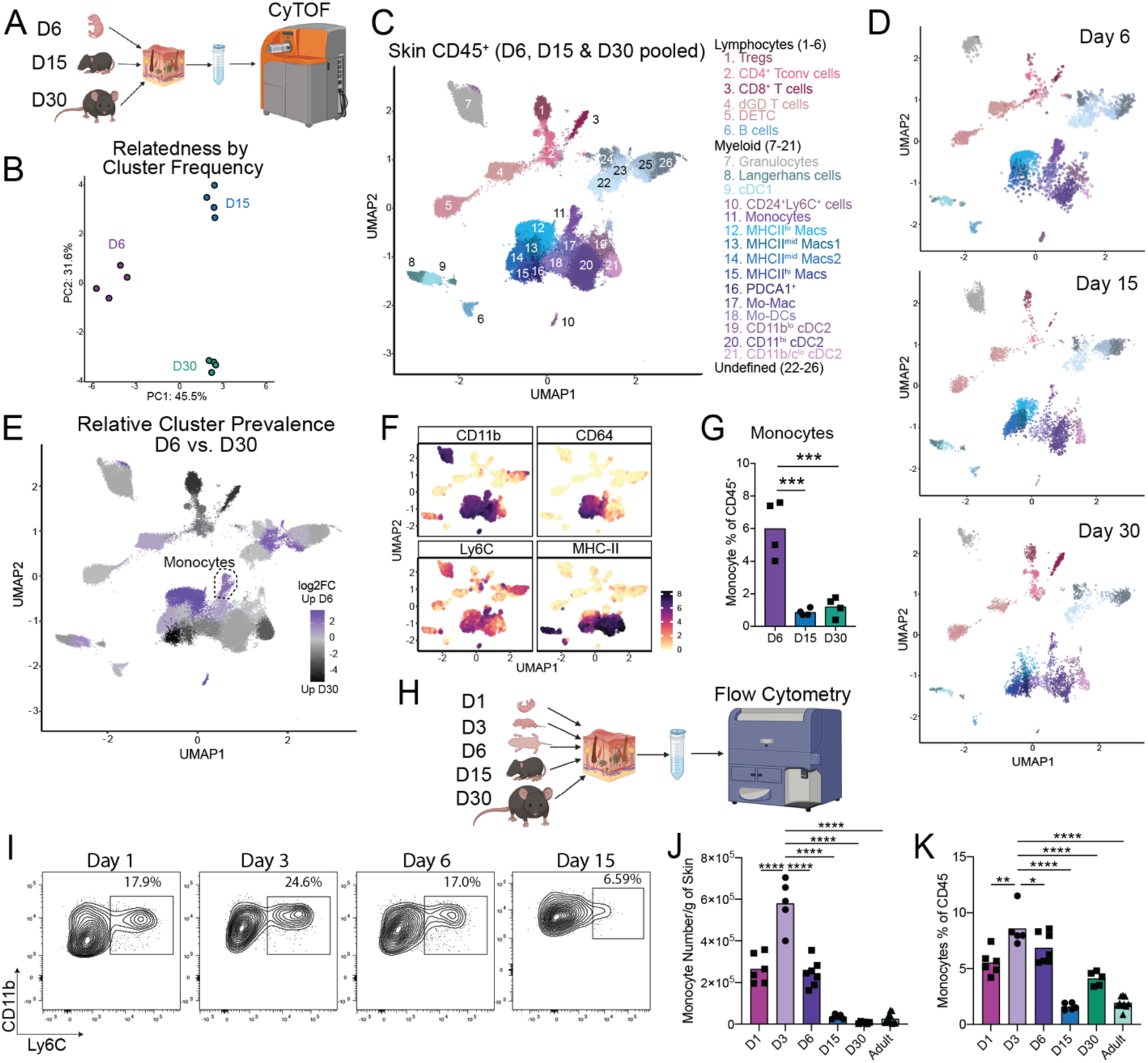
Classical monocytes rapidly accumulate in neonatal skin. (A) Skin tissue from mice at the postnatal Day 6, Day 15 and Day 30 ages was harvested and analyzed by Mass Cytometry (CyTOF). (B) Principal component analysis (PCA) plot demonstrating the distribution of the cutaneous CD45^+^ cellular compartment based on cluster frequency. Each dot represents a single mouse at the indicated age. (C) Pooled UMAP plot demonstrating clusters of CD45^+^ immune cells in the skin of mice at Day 6, 15 and 30. (D) Individual UMAPs demonstrating immune cell clusters in the skin of Day 6, 15 and 30 mice. (E) UMAP demonstrating clusters of immune cells enriched in the skin of Day 6 (purple) or Day 30 (black) mice. (F) UMAPs demonstrating expression profile of indicated markers to identify monocyte cluster. (G) Enumeration of monocytes as a percent of the total immune cell compartment of the skin at the indicated time points. (H) Back skin from mice at postnatal day 1, 3, 6, 15 & 30 and >8-week-old adult mice was analyzed by Flow Cytometry. (I) Representative flow cytometry plots of skin monocytes at the indicated ages. (J-K) Enumeration of skin monocytes by flow cytometry as (J) total numbers per gram of skin and (K) percentage of CD45^+^ cells. For CyTOF experiment n=4 mice were used at each time point, for flow cytometry assays >4 mice were used at each time point. Data presented in I-K is one representative of at least two independent experiments. Statistical significance was determined by a One-Way ANOVA where *** is p<0.001 and **** is p<0.0001.

Examination of lymphoid populations revealed stable percentages of Dendritic Epidermal T cells (DETCs) between postnatal day 6 and day 30, consistent with their known prenatal skin seeding^18^ (Fig. S1C). Likewise, this dataset confirmed our prior flow cytometry studies demonstrating rapid accumulation of regulatory T cells (Tregs) in skin during the first two weeks of postnatal life, with a subsequent increase in conventional CD4^+^ and CD8^+^ T cells^5,19^ (Fig. S1D-G).

Examination of myeloid cells also revealed age-dependent shifts in relative cluster abundance. Consistent with recent work^11^, type 1 classical dendritic cells (cDC1) were somewhat enriched at the immediate postnatal window, in contrast prevalence of their type 2 counterparts (cDC2) increased at later time points (Fig. S1H). While most other myeloid clusters and the total proportion of myeloid cells had their peak relative abundance at day 15 or later (Fig. 1D-E, Fig. S1G), we noted three clusters that were highly enriched at postnatal day 6. We identified the first of these as MHC-II^lo^ macrophages (cluster 12), which were replaced progressively at days 15 and 30 of life by MHC-II^mid^ and MHC-II^hi^ macrophages, consistent with a gradual process of maturation and activation in the tissue^20^ (Fig. S1I-K). The second was a small CD11b^+^ MHCII^neg^ population (cluster 10) that was intermediate for CD24 and Ly6C, and hard to definitively classify (Fig. S1L). The third expressed markers consistent with a monocyte identity, i.e. Ly6C^hi^ CD11b^hi^ CD24^lo^ CD64^lo^ MHCII^neg^ (cluster 11) (Figs. 1F, S1B). These monocytes demonstrated a high day 6 abundance (cumulatively ∼6% of all CD45^+^ cells), followed by a drop off in proportion/density by the day 15 time point (Fig. 1G). Characterization of these neonatal skin monocytes by flow cytometry confirmed their high expression of CCR2 and Ly6C, further identifying them as classical monocytes (Fig. S1M).

Classical monocytes are typically considered pro-inflammatory cells studied in the context of tissue infection or injury^12^. Thus, we were intrigued to see them enriched in neonatal skin, which has a pro-tolerogenic immune tone towards commensal microbes and other antigens^1,19^. To further define the timing of skin monocyte seeding, we performed longitudinal enumeration of skin monocytes by flow cytometry from postnatal day 1 to 30 (Fig 1H). Although skin monocytes were present in 1-day old pups, their percentage and number dramatically increased by postnatal day 3 (Fig. 1I-K). Consistent with our CyTOF data, monocyte numbers remained substantially higher at day 6 before tapering off at later timepoints, likely reflecting their known propensity to downregulate Ly6C and gain MHC-II expression as part of their differentiation into monocyte-derived dendritic cells (Mo-DCs)^21^. In contrast, there were no substantial age-dependent differences in the proportion of monocytes in the spleen over this period (Fig. S1N). Cumulatively, these data demonstrate that the myeloid immune landscape in the skin changes with age and that a population of classical monocytes (neoMono) accumulate in high density and proportion in the early life window.

### Commensal microbes facilitate early skin accumulation of monocytes

Given the postnatal timing of neoMono accumulation in skin, we hypothesized that recognition of microbial ligands might provide the signals necessary to drive their recruitment. To examine this, we performed flow cytometry on skin from day 6 germ-free (GF) and specific-pathogen-free (SPF) pups, which revealed significantly higher neoMono numbers and percentages in the back skin of SPF pups as compared to GF pups (Fig. 2A-B). To further assess the requirement for commensal colonization in neoMono skin accumulation we subjected pregnant SPF dams to a cocktail of antibiotics in the drinking water along with frequent cage changes to minimize the microbial load in the bedding. Pups born in these “reduced microbial burden” cages had significantly lower percentages and numbers of skin neoMono on postnatal day 6 (Fig. 2C-D). To further demonstrate the sufficiency of commensal microbes in promoting neoMono accumulation, we colonized the skin of germ-free pups with *Staphylococcus epidermidis* within the first 2 days of life. Exposure to this single skin commensal bacterium significantly increased the percentage and number of neoMono in the skin at postnatal day 6 (Fig. 2E-F).

**Figure 2:**
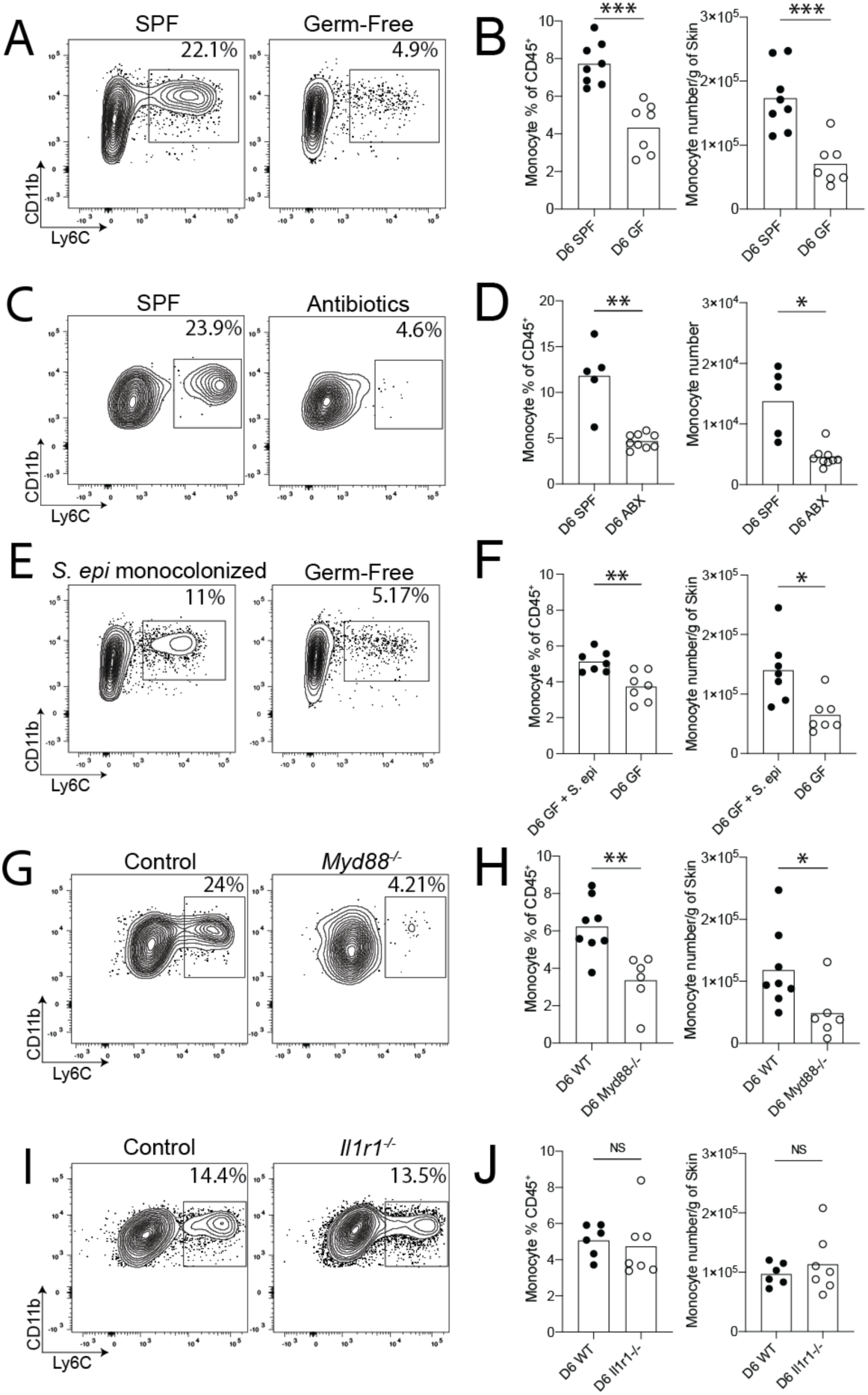
Commensal microbiota facilitate early skin accumulation of monocytes. Neonatal skin from mice at the postnatal Day 6 age was harvested and analyzed for monocytes using flow cytometry. Representative flow cytometry plots and quantification of monocyte numbers from (A-B) Specific Pathogen Free (SPF) control and Germ-Free (GF) pups, (C-D) SPF control pups and pups born to antibiotic treated dams in cleaner environments, (E-F) GF pups colonized on postnatal Day 2 of life with *S. epidermidis* and age-matched GF controls, (G-H) Control and *Myd88^-/-^* pups, (I-J) Control and *Il1r1^-/-^* pups. Data from (A-B and G-H) are pooled from two experiments where each point represents a single mouse with a pooled n≥5 mice per group, per condition. Data from (C-D, E-F and I-J) are one representative of at least two independent experiments with n≥5 mice per group, per condition. Statistical significance was determined by a two-tailed Student’s t test where * is p<0.05, ** is p<0.01 and *** is p<0.001

Recognizing that commensal microbes can tune levels of alarmin signaling in barrier tissues and that this might serve as a signal for cell recruitment, we next assessed neoMono numbers in the back skin of mice lacking either Myeloid differentiation primary response 88 (*Myd88*^-/-^) or interleukin-1 receptor 1 (*Il1r1*^-/-^). At postnatal day 6, *Myd88*^-/-^ (Fig. 2G-H) but not *Il1r1*^-/-^ (Fig. 2I-J) mice demonstrated significantly lower numbers of skin monocytes. Taken together these data indicate that neoMono skin accumulation is dependent on Myd88 but not IL-1R1 signaling, suggesting that toll-like receptor detection of microbial ligands upstream may be involved.

### Neonatal monocyte depletion via anti-Gr1 antibody affects type 17 signature of skin T cells

To uncover the biological significance of neoMono skin accumulation, we developed a temporal antibody-mediated depletion strategy in the first two weeks of life. By delivering the anti-Gr-1 antibody, which targets cells expressing Ly6G or Ly6C, intraperitoneally to neonatal mice every other day between day 4 and 14 of life, we achieved substantial depletion of skin neoMono as confirmed by flow cytometry on postnatal day 15 (Fig. 3A-C). To gain a broader understanding of the impact of neoMono depletion on the composition and function of the cutaneous immune compartment, we performed single cell RNA-sequencing (scRNA-seq) on CD45^+^ cells sorted from the back skin of anti-Gr-1-treated (NeoΔMono) versus isotype-treated (Control) mice on postnatal day 15 (Fig. 3A). Our analysis showed 20 clearly distinguishable clusters (Fig. 3D) which, based on their expression of key markers (Fig. S2A-B) could be primarily categorized as lymphoid cells or myeloid cells with the exception of one cluster defined primarily by their proliferative status and a small number of contaminating keratinocytes (KC) (Fig. 3E).

**Figure 3:**
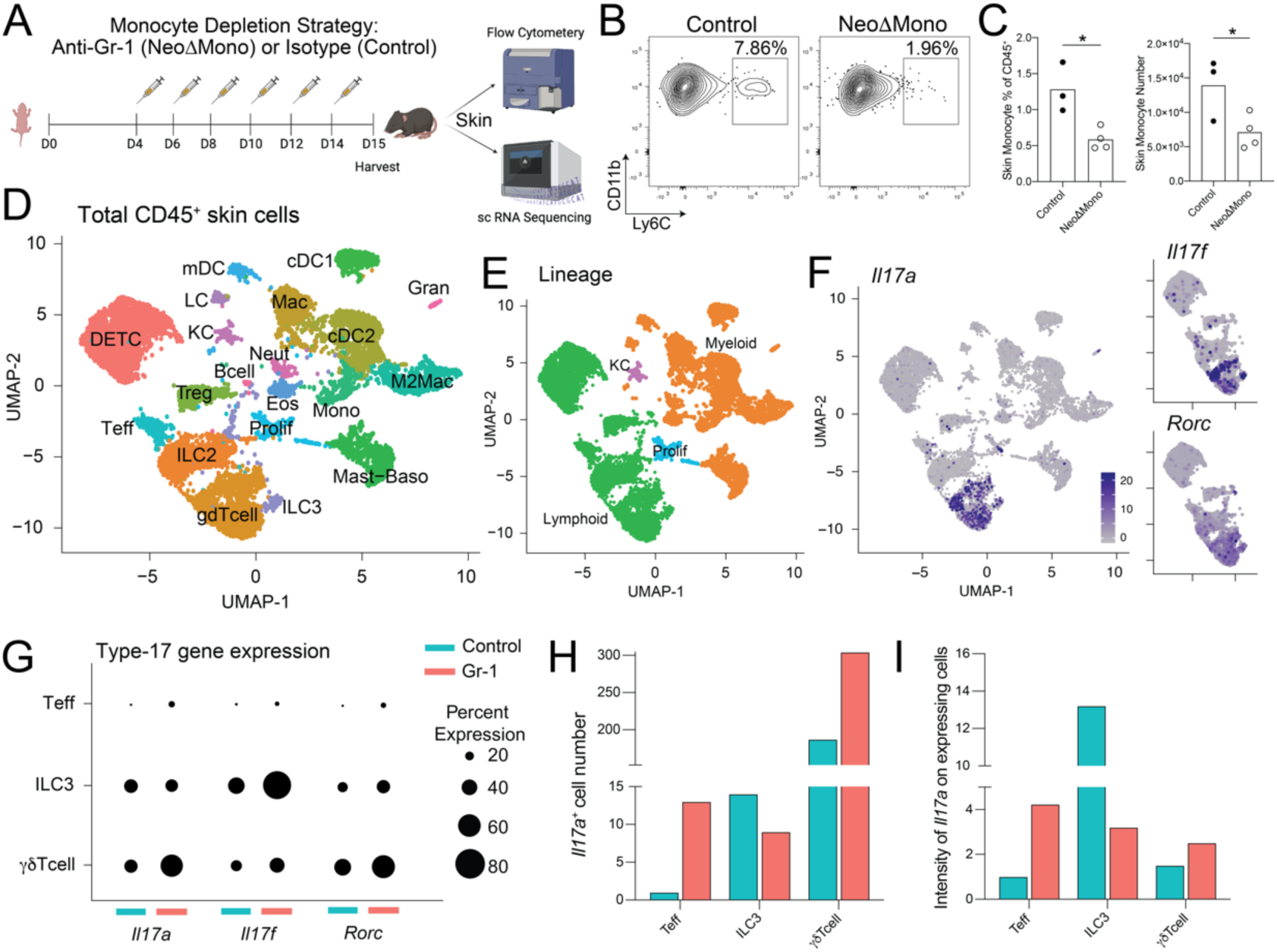
Neonatal monocyte depletion affects type 17 signature of skin T cells. (A) Monocytes were depleted in neonatal mice by treatment with the anti-Gr-1 antibody (NeoΔMono mice) and controls were treated with isotype antibody. Skin from these mice was then processed either for flow cytometry or scRNA-seq. (B-C) Representative flow plots, with percent of parent gate shown, and quantification of monocytes, as % of CD45^+^ and absolute numbers, from the skin of NeoΔMono mice and controls at postnatal Day 15. (D-E) UMAP from scRNA-seq analyses demonstrating clusters of immune cell populations in the skin of NeoΔMono and control mice. (F) UMAPs demonstrating expression profiles of *Il17a*, *Il17f* and *Rorc* in skin lymphoid clusters of NeoΔMono and control mice, for the latter two only lymphoid clusters are shown. (G) Dot plot demonstrating the percent of cells within the cutaneous Teff, ILC3 and γδ T cell clusters that express *Il17a*, *Il17f* and *Rorc* in NeoΔMono and control mice. (H-I) Quantification of the number of *Il17a* expressing cells and intensity of *Il17a* expression in Teff, ILC3 and γδ T cell clusters. For C, statistical significance was determined by a two-tailed Student’s t test where * is p<0.05. All scRNA-seq analyses were performed post normalizing clusters across both treatment groups.

Examination of relative cluster abundance (Fig. S2C-E) did show a substantial relative reduction in skin monocytes in NeoΔMono versus control skin. Confirmatory flow cytometry studies confirmed anti-Gr-1 depletion of a small population of skin neutrophils, as anticipated based on their Ly6G expression (Fig. S2F). Other myeloid cell populations such as dendritic cells and macrophages were not significantly affected by neonatal anti-Gr-1 treatment at this time point (Fig. S2G-H).

Our sequencing and flow cytometry data did not reveal substantial differences in the relative proportion or number of effector (i.e. Foxp3^neg^) CD4^+^ T cells (Teff), Tregs, or γδ T cells in postnatal day15 NeoΔMono versus control skin (Figs. S2E and S2I). However, further examination of transcripts revealed a shift towards a heightened type 17 signature among CD4^+^ Teff and dermal γδ T cells, as evidenced by an increased percentage of cells in each cluster expressing *Il17a, Il17f* or *Rorc* (Fig. 3F-G). For *Il17a*, in particular, the total number of *Il17a-* expressing Teff and γδ T cells and the intensity of *Il17a* expression on these cells was increased in NeoΔMono mice (Fig. 3H-I). In contrast, type 3 innate lymphoid cells (ILC3) in NeoΔMono mice showed an overall reduced percentage and intensity of *Il17a* expression (Fig. 3H-I). These data suggest a possible regulatory effect of neoMono on lymphocyte production of the key barrier cytokine IL-17.

### Transient early life monocyte depletion leads to sustained enrichment of IL-17A producing skin CD4^+^ T cells

To further explore this shift in skin lymphocyte IL-17A production following early life anti-Gr-1 treatment, we performed flow cytometry experiments on NeoΔMono or isotype antibody-treated mice (Fig 4A). Examination of the back skin on day 21 confirmed an increase in the percentage and number of IL-17A producing CD4^+^ Teff (Figs. 4B and S3A) and dermal γδ T cells (Figs. 4C and S3A). This was also reflected in the corresponding SDLN populations (Fig. 4D-E). As we have previously shown the importance of neonatal skin Tregs in helping to establish tissue immune homeostasis, we examined the skin CD4^+^ compartment in anti-Gr-1 treated mice which did not reveal significant differences in either the percentage or number of skin Tregs (Fig. 4F) or Teffs (Fig. 4G).

**Figure 4:**
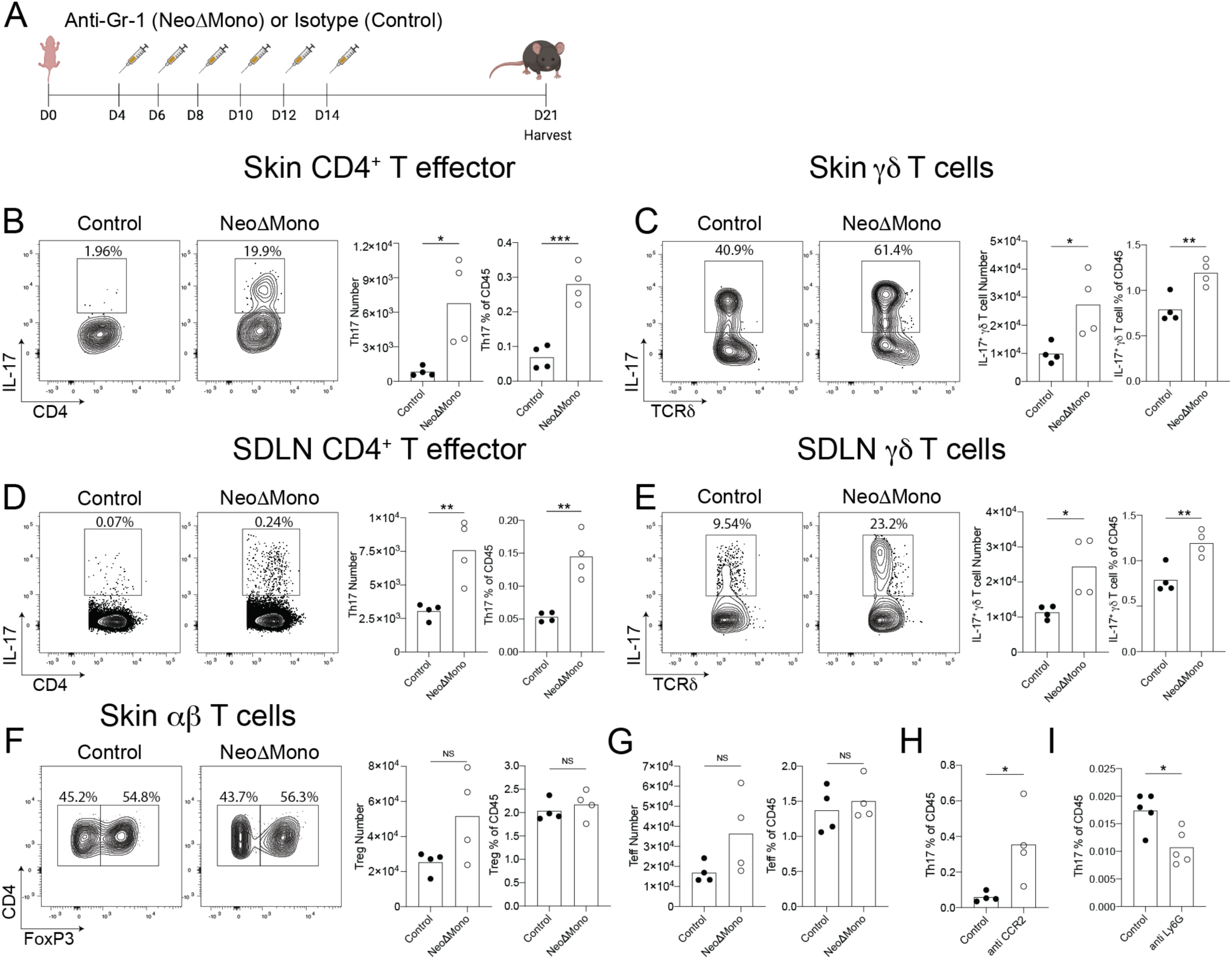
Transient depletion of monocytes in early life leads to sustained type 17 inflammatory responses from cutaneous T lymphocytes. (A) Monocytes were depleted in neonatal mice by treatment with the anti-Gr-1 antibody (NeoΔMono mice) and controls were treated with isotype antibody. Skin from these mice were then processed for flow cytometry at postnatal Day 21. (B & D) Representative flow cytometry plots and quantification of numbers of CD4^+^ T effector cells producing IL-17 in the skin (B) and skin draining lymph nodes (D). (C & E) Representative flow cytometry plots and quantification of numbers of γδ T cells producing IL-17 in the skin (C) and skin draining lymph nodes (E). (F-G) Representative flow cytometry plots and quantification of numbers of CD4^+^ T regulatory (F) and effector (G) cells in the skin. (H-I) Quantification of numbers of CD4^+^ T effector cells producing IL-17 in the skin of mice treated with anti-CCR2 antibody (H) or anti-Ly6G antibody (I) according to the same schedule as in (A). Data in A-G are one representative of at least three independent experiments, H-I are one representative of at least two independent experiments. Statistical significance was determined by a two-tailed Student’s t test where * is p<0.05, ** is p<0.01 and *** is p<0.001

To understand timing of this observed type 17 immune shift, we performed flow cytometry on analogously treated mice on day 15, just after anti-Gr-1 treatment was completed, and 3.5 weeks later on day 40 of life. The day 15 data paralleled our scRNA-seq dataset from the same timepoint showing increases in IL-17A^+^ skin Teff and dermal γδ T cells (Fig. S3B-C). By day 40, however, it was only the increased skin IL-17A^+^ skin Teff (Th17) phenotype that persisted in NeoΔMono mice, whereas IL-17A production by dermal γδ T cell had normalized (Fig. S3B-C).

To further verify that the type 17 phenotype was in fact due to the absence of neoMono, we performed analogous experiments delivering an anti-Ccr2 antibody between postnatal day 4 and 14. This alternative monocyte-targeting approach also led to an increase in the percentage and number of skin Th17 cells on postnatal day 21 (Fig. 4H, S3D). As anti-Gr-1 also depletes neutrophils, we performed neonatal delivery of anti-Ly6G antibody to specifically target neutrophils but not monocytes, which did not recapitulate the increased type 17 phenotype (Fig. 4I, S3E). We measured monocyte numbers on postnatal Day 21 and 30 after anti-Gr-1 treatment (Fig S3F-G). No significant differences were noted between NeoΔMono and control groups, thus validating a recovery of monocytes at the time points used to assess type 17 responses. To ascertain if the type 17 phenotype was specific to the anti-Gr-1 treatment in neonatal life or an age independent effect, we performed the anti-Gr-1 treatment in adult mice (Fig. S3H). This did not reveal any significant shifts in IL-17A production by either skin CD4^+^

Teff (Fig. S3I) or dermal ψ8 T cells (Fig. S3J), indicating this response is specific to targeting of monocytes in the early life window. Collectively these data suggest that specifically eliminating the observed early life accumulation of neoMono leads to an increase in IL-17A production by skin T cells.

### Commensal skin microbes contribute to elevated type 17 T cells in NeoΔMono mice

Myeloid cells are an important population interacting with skin microbes^11,22^ and type 17 responses are a typical flavor of immunity elicited by both commensal and pathogenic skin bacteria^23–25^. We thus hypothesized that the depletion of monocytes might lead to an altered skin microbial community. To test this, we sampled the skin of NeoΔMono and co-housed control littermates on postnatal day 15, one day after the last anti-Gr1 treatment, and performed 16S rRNA sequencing to assess bacterial community composition. Measurement of beta diversity revealed significant differences between NeoΔMono and control littermates (Fig. 5A). Further analysis revealed that major drivers of this difference included a decreased relative abundance of *Staphylococcus* and an increased relative abundance of *Bacteroides* on the skin of NeoΔMono mice as compared to littermate controls (Fig. 5B-C). Thus, even before peak differences in type 17 inflammation are seen in NeoΔMono mice, there is a measurable shift in the composition of the skin microflora.

**Figure 5:**
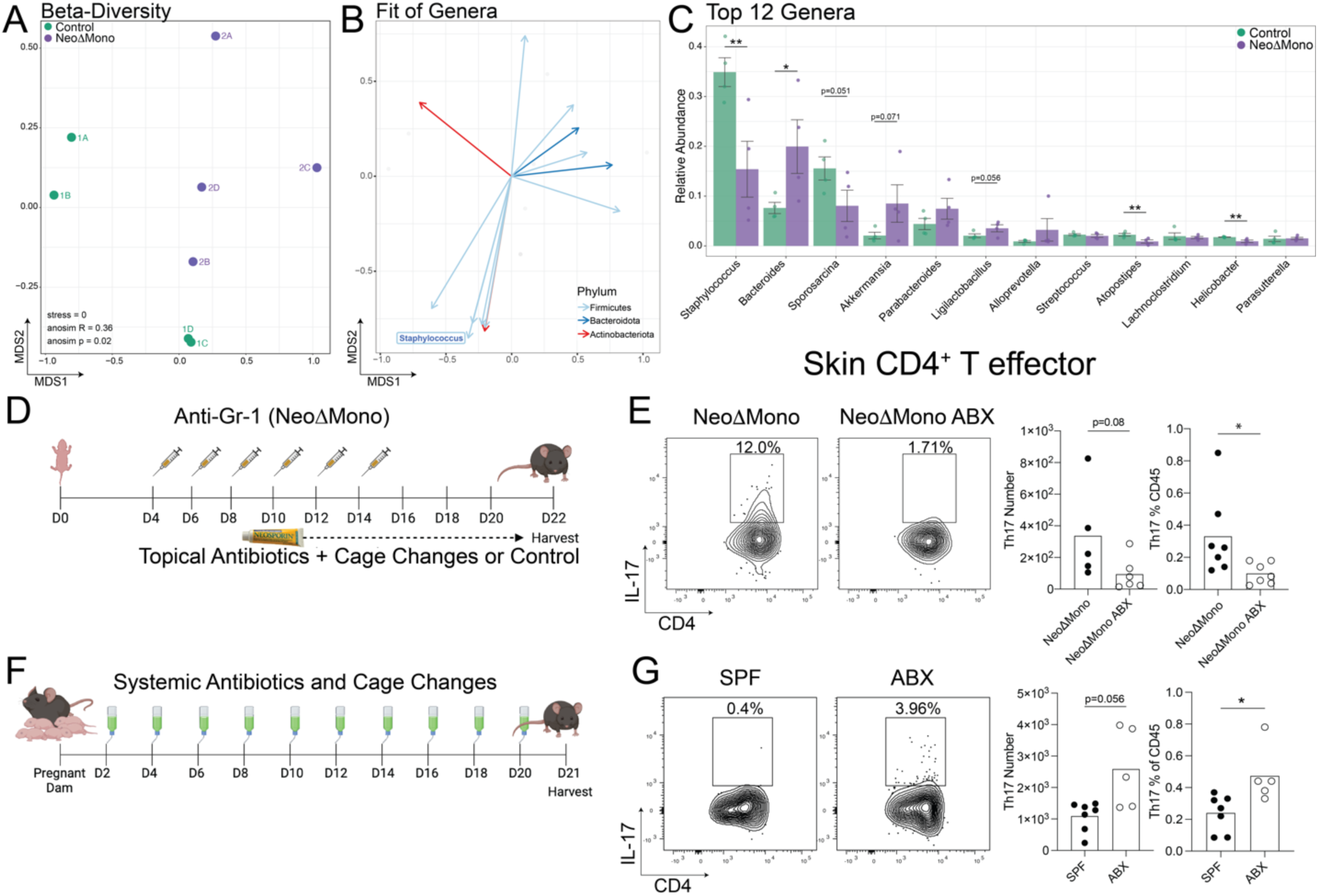
Commensal microbes contribute to elevated type 17 response in NeoΔMono mice. Skin swabs were taken from NeoΔMono mice and littermate controls on postnatal day 15 and processed for 16S sequencing. (A) Non-metric multidimensional scaling (NMDS) plot calculated using Bray-Curtis dissimilarities of Amplicon Sequence Variants (ASVs) demonstrating the differences in beta-diversity of bacterial communities in day 15 skin between NeoΔMono mice and littermate controls. (B) Environmental fit of ASVs aggregated at the genus level, excluding unassigned genera, driving spatial distribution of points in the NMDS in A. (C) Bar plot showing mean and standard error of the relative abundances of the top 12 most abundant genera, in descending rank-order, in the skin of either group of mice. (D) NeoΔMono mice were treated with topical Neosporin and frequent cage changes while controls were treated with vehicle and subjected to mock cage changes. Skin was then processed for flow cytometry on postnatal Day 22. (E) Representative flow cytometry plots and quantification of numbers of IL-17-producing CD4^+^ T effector cells in ear skin. (F) Pregnant dams were treated with a cocktail of antibiotics in the drinking water along with frequent cage changes until pups were 21 days old. (G) Representative flow cytometry plots and quantification of numbers of IL-17-producing CD4^+^ T effector cells in back skin. Data in E are pooled from two of three experiments, data in F is from one of two independent experiments. Analysis of Similarity (*anosim*; vegan R package) was used to test for differences in the community structures between both treatment groups with statistical significance displayed in plot for (A). For (B) each arrow shows a correlation with the relative abundance and coordinate space for genera with p</=0.05, R>/=0.1. Statistical significance in C, E, and G was determined by a two-tailed Student’s t test where * is p<0.05, and ** is p<0.01.

We next sought to test if the heightened type 17 response in NeoΔMono was facilitated by the presence of skin bacteria. To test this, we performed early life anti-Gr-1 treatment in tandem with an established protocol to broadly deplete skin bacteria^26,27^, namely topical application of Neosporin to the ears plus frequent cage changes (Fig. 5D). Examination of skin T cells by flow cytometry on day 22 showed a reduction in the percentage and number of IL-17A producing skin CD4^+^ Teff (Fig. 5E) and dermal ψ8 T cells (Fig. S5A), indicating that the presence of neoMono helps to limit the homeostatic immune response to skin bacteria. Next, we evaluated the impact of extended antibiotic exposure from birth on the generation of altered type 17 immune responses in the skin. Pregnant dams were given antibiotics in the drinking water and subjected to frequent cage changes until their pups were 21 days old (Fig. 5F). These interventions aimed at lowering overall microbial burden led to significantly reduced numbers of monocytes in the skin even at postnatal day 21 (Fig. S4C) and increased numbers of IL-17 producing CD4^+^T cells and ψ8 T cells (Fig. 5G and S4B). These data indicate that early-life disruptions in microbial community composition can contribute to aberrant type 17 inflammation from T cells in the skin at homeostasis.

### NeoΔMono mice demonstrate a heightened inflammatory response to imiquimod-driven psoriasiform inflammation

To elucidate the physiological impact of the dysregulated type 17 inflammatory response in anti-Gr-1 treated NeoΔMono mice, we utilized a well-established murine model of IL-17 driven psoriasiform skin inflammation. Ear skin of 4 week-old NeoΔMono and control mice were treated topically every day for 5 days with 5% imiquimod cream (Fig. 6A). Ear thickness was measured daily using digital calipers as an indicator of gross inflammation and ear skin and draining lymph nodes were harvested at day 6 for flow cytometric analyses. NeoΔMono mice demonstrated greater ear swelling as early as day 3 of treatment, which became even more substantial by day 5 (Fig. 6B). This ear swelling was accompanied by increased histologic markers of skin inflammation, including a thickened stratum corneum with dysregulated corneocyte differentiation (parakeratosis), accentuated hair follicle plugging and a substantial inflammatory infiltrate in the dermis and subcutis (Fig. 6C-D). Enumeration by flow cytometry (Fig. S5A) revealed a significantly higher neutrophil percentage in the ears (Fig. 6E) and ear-draining lymph nodes (Fig. 6G) of NeoΔMono mice. This paralleled an increase in the percentage and number of Th17 cells in NeoΔMono ear skin (Fig. 6F), which was largely recapitulated in the ear-draining lymph nodes (Fig. 6H, S5B-C). Notably, even though inflammation in this model is known to have a strong contribution from IL-17A-producing dermal γδ T cells, we did not note significant differences in their percentages or numbers between groups (Fig. S5B-C), consistent with our observation that spontaneous IL-17A-production by dermal γδ T cells returns to baseline levels by day 40 (Fig. S3C). Taken together these data suggest that neonatal depletion of monocytes can render mice more susceptible to an IL-17 driven psoriasiform skin inflammation.

**Figure 6:**
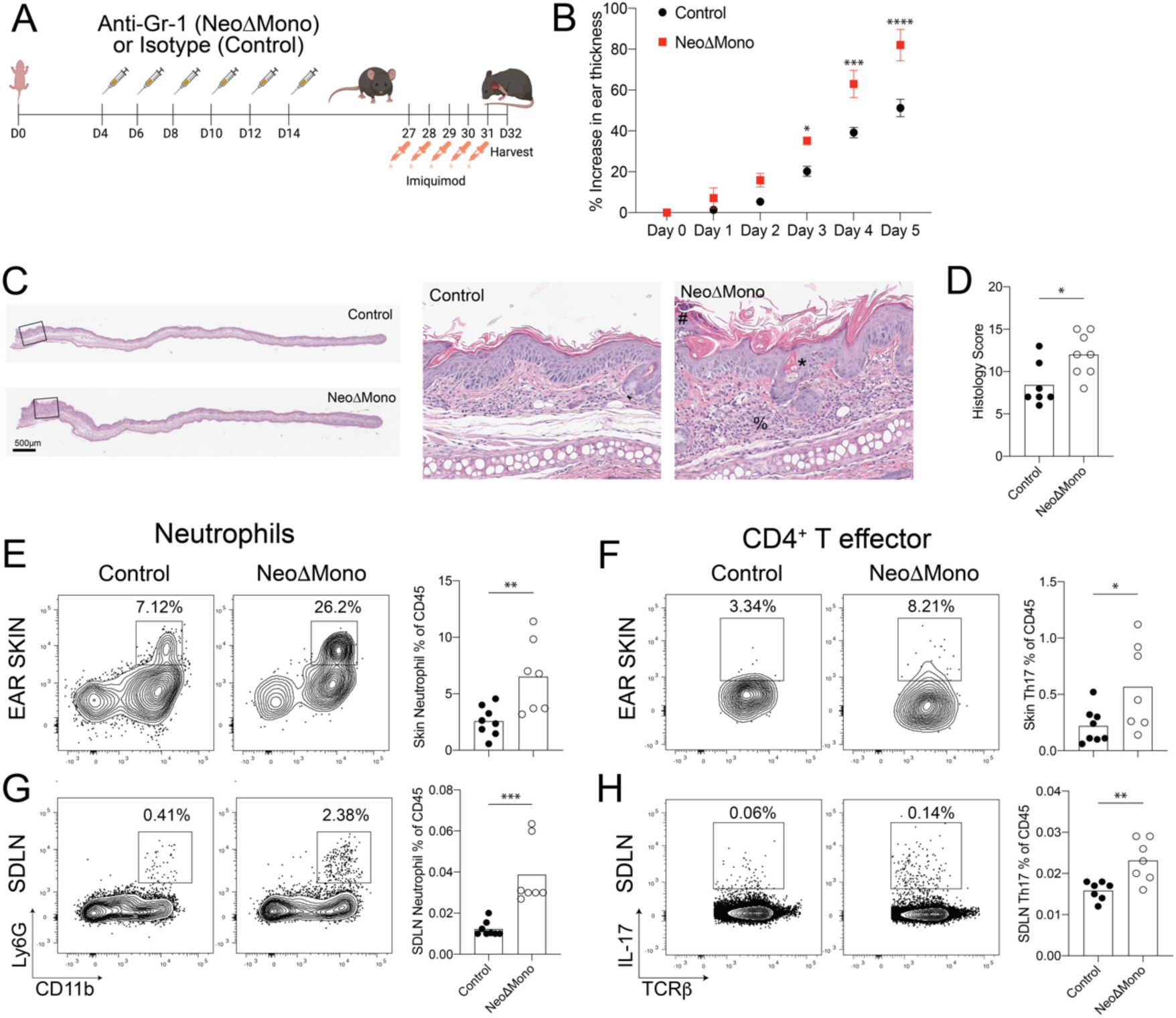
NeoΔMono mice demonstrate a heightened inflammatory response to imiquimod-driven psoriasiform inflammation. (A) Monocytes were depleted in neonatal mice by treatment with the anti-Gr-1 antibody (NeoΔMono mice) and controls were treated with isotype antibody. The ears of these mice were treated daily for five days with 5% Imiquimod cream then one day later ear skin and lymph nodes were harvested. (B) Quantification of relative increase in ear thickness as measured with digital calipers during imiquimod treatment in NeoΔMono versus control mice. (C-D) Representative images of ear skin sections stained with H&E with zoomed inset (C) and quantification of histopathology scoring (D) between groups. # represents parakeratosis (neutrophils in the stratum corneum), * represents hyperkeratosis and plugging and % represents dermal and subcutaneous inflammatory infiltrate. (E-F) Representative flow cytometry plots and quantification of neutrophils (E) and CD4^+^ Th17 cells (F) in the skin. (G-H) Representative flow cytometry plots and quantification of neutrophils (G) and CD4^+^ Th17 cells (H) in the skin draining lymph nodes. Data in B is from one of three independent experiments. Data in E-H is data pooled from two independent experiments. Statistical significance in B was determined by a two-way ANOVA with Sidak’s multiple comparisons test where * is p<0.05, *** is p<0.001 and **** is p<0.0001. Statistical significance in E-H was determined using a two-tailed Student’s t test where * is p<0.05, ** is p<0.01 and *** is p<0.001.

We performed similar studies in adult mice where monocytes were depleted at 8 weeks of age or older using the anti-Gr-1 antibody (AdultΔMono), then two weeks later ear skin was treated topically with imiquimod for 5 days (Fig. S5D). As predicted from the lack of spontaneously increased type 17 inflammation following adult depletion of monocytes, even after imiquimod no significant differences were noted in ear thickness, numbers of neutrophils, Th17 or IL-17 producing ψ8 T cells in either ear skin or draining lymph nodes (Fig. S5E-G). To test the durability of the heightened inflammatory response to imiquimod following neoMono depletion, we challenged NeoΔMono mice at 8 weeks of age, a full 6 weeks after the last anti-Gr-1 treatment (Fig. S5H). At this later time point, no statistically significant differences in the measured parameters were noted between imiquimod-treated NeoΔMono on control mice (Fig. S5I-K). Collectively, these data illustrate that neonatal monocytes play a key role in regulating type 17 responses and susceptibility to psoriasiform skin inflammation for a sustained but not indefinite period.

### Neonatal myeloid cells preferentially express IL-1 pathway repressors

We next set out to decipher the mechanism driving the dysregulated type 17 response in the skin of NeoΔMono mice, with a particular focus on CD4^+^ Teff as the most persistently affected population. To achieve this, we returned to our scRNA-seq data from D15 control skin to gain deeper insights into the transcriptional signatures of myeloid cell populations in neonatal skin, especially the dominant monocyte and monocyte-derived populations at this time point. Using unsupervised clustering and top differentially expressed genes per cluster we identified a cluster with overlapping features of monocytes (S100a8, Il1b, Tlr2, Plac8^28,29^ and macrophages (Msr1, Tgfbi, Fcgr1^28–31^ (i.e. mono-Mac) (Fig. 7A, S6A). This was suggestive of monocytes starting to undergo differentiation within the tissue^28,32^, consistent with this being a day 15 time point, almost two weeks after the peak neoMono accumulation (Fig. 1J). In addition to these transitioning cell types we identified various subsets of macrophages, including (Mature Macs) (H2-Aa, H2-Ab1, H2-Eb1, H2-DMb1, H2-DMb2^28,32^), and tissue-resident macrophage (TR-M) (Selenop, and Folr2)^33^. Other clusters included cDC2, migratory DCs (mDCs), Langerhans cells (LC), various granulocytes and some contaminating keratinocytes (Fig. 7A, 6SA). We then performed gene set enrichment analysis which revealed an increase in IL-1 pathway gene expression among various myeloid subsets. In particular, the negative regulators of this pathway, namely the IL-1 receptor antagonist IL-1Ra (*Il1rn*) and the decoy receptor IL-1R2 (*Il1r2*), were expressed at very high levels in several myeloid cell types, including the Mono-Mac cluster (Fig. 7B-C, Fig. S6B-C). Further gene set enrichment analysis revealed that this Mono-Mac cluster, representing cells occupying an intermediate state between early monocytes and mature macrophages, was enriched for IL-1 pathway genes when compared to other neonatal skin macrophage and dendritic cell subsets (Fig. 7D, S6D). To determine if IL-1 pathway repressors are also expressed in skin monocytes just entering this tissue, we performed qPCR analysis for *Il1rn* and *Il1r2* on FACS-sorted monocytes and macrophages isolated from the skin of 4 day-old pups and adult mice. This showed higher *Il1rn* and *Il1r2* transcript levels in monocytes versus mature macrophages in neonatal skin (Fig. 7E, S6E, gating S1M). Additionally, it revealed that this heightened IL-1 pathway repressor expression was particularly enriched in day 4 skin monocytes as compared to those in day 30 skin (Fig. 7E, S6E). Collectively, these data indicate that during early life in the skin, monocytes are enriched in molecules poised to regulate IL-1 signaling, a pathway known to facilitate IL-17A production by CD4^+^ T lymphocytes^26^.

**Figure 7:**
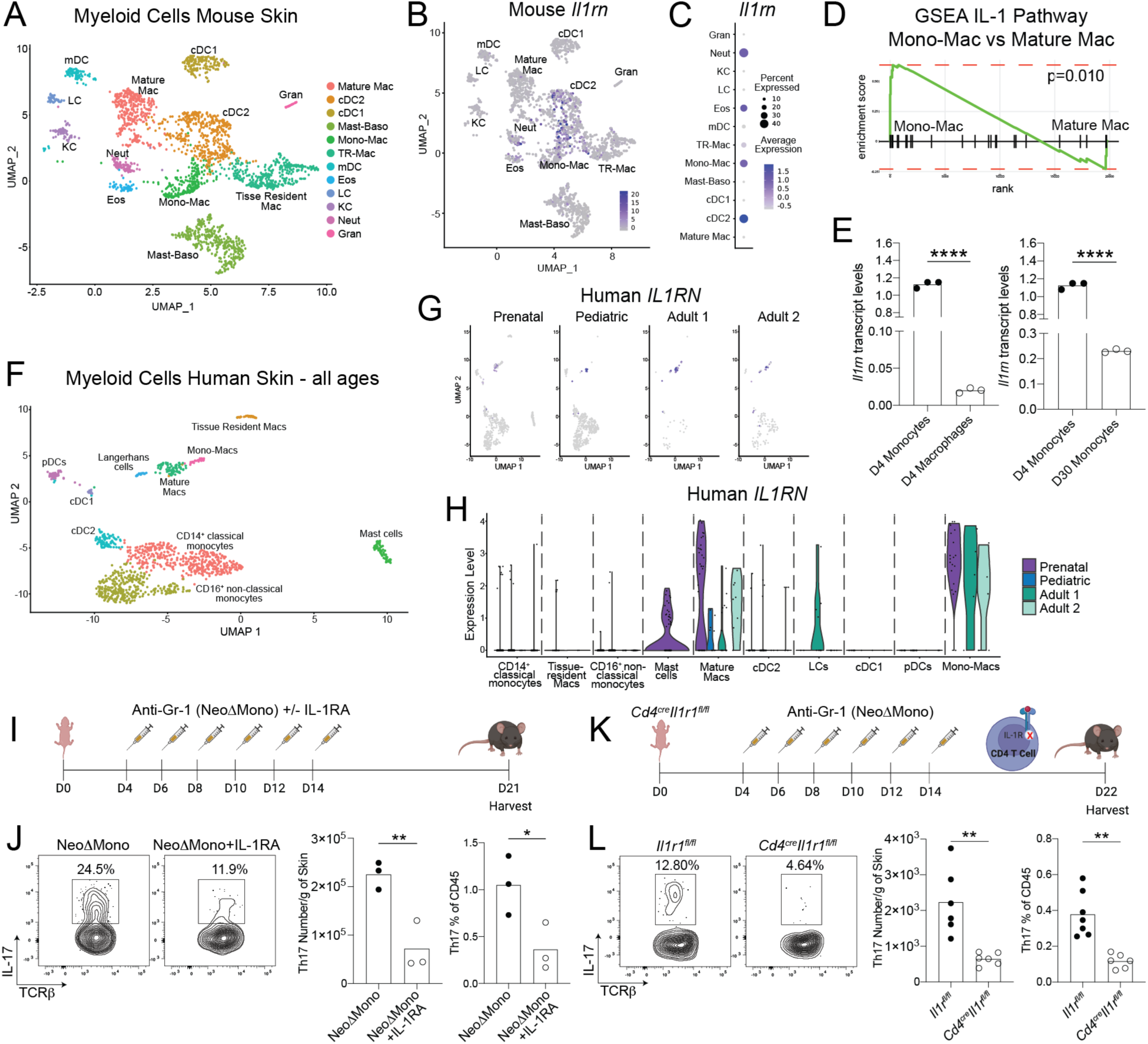
Neonatal monocytes modulate IL-1 signaling to regulate cutaneous type 17 inflammation. (A) UMAP of myeloid cluster from scRNA-seq analysis of postnatal Day 15 mouse skin. (B) Intensity plot showing *Il1rn* expression across murine skin myeloid clusters. (C) Quantification of *Il1rn* expression by normalized average expression and percentage of expressing cells for individual myeloid clusters. (D) Gene Set Enrichment Analyses (GSEA) for IL-1 pathway genes comparing postnatal Day 15 skin mono-mac and mature mac clusters. (E) qPCR analysis of *Il1rn* transcript levels comparing monocytes versus macrophages sorted from postnatal Day 4 skin and monocytes sorted from postnatal Day 4 versus postnatal Day 30 skin. (F) UMAP of myeloid clusters from scRNA-seq analysis of human skin – combining cells from all four donors. (G-H) Intensity and violin plots showing *IL1RN* expression across human skin myeloid clusters by age. (I) NeoΔMono mice were either treated with recombinant IL-1Ra or with vehicle. Skin was then processed for flow cytometry on postnatal Day 21. (J) Representative flow cytometry plots and quantification of numbers of CD4^+^ T effector cells producing IL-17A in the skin. (K) Monocytes were depleted in neonatal *Cd4^cre^Il1r1^fl/fl^* or control *Il1r1^fl/fl^* mice by treatment with the anti-Gr-1 antibody (NeoΔMono mice). Skin was then processed for flow cytometry on postnatal Day 21. (L) Representative flow cytometry plots and quantification of numbers of CD4^+^ T effector cells producing IL-17A in the skin. Statistical significance was determined using a two-tailed Student’s t test where * is p<0.05, ** is p<0.01 and **** is p<0.0001. Data in J is representative data from one of three independent experiments and data in L is pooled from two independent experiments.

To determine if early life expression of IL-1 pathway repressors is conserved in skin myeloid cells across species, we next performed scRNA-seq on immune cells from human skin, including on prenatal, one pediatric (age 16) and two adult samples (ages 33 and 36). Performing clustering on cells from all samples, we were able to clearly identify expected populations of lymphoid and myeloid immune cells (Fig. S6F). Of note, looking at cluster frequencies across each sample there was a clear trend toward a larger proportion of myeloid cells in our younger samples versus T cells, as we observed in mice (Fig. S1G, S6G). Looking more closely at the myeloid cell types we were able to identify clusters, analogous to most of those observed in mice, expressing markers consistent with transitioning monocytes (Mono-Macs) (TLR2, MSR1, PLAC8, TGFBI and FCGR1A^28–31,34^), mature macrophages (HLA-DQB1, HLA-DRA, HLA-DRB1, HLA-DMA and HLA-DOA^28,32^), TR-Ms (LYVE1, MRC1, FOLR2, FOS, MAF^33^), LCs (CD207), cDC2 (CLEC10A^29,34,35^), cDC1 (CLEC9A^36^, pDCs (LILRA4^37^), and mast cells (TPSB2^29^). (Fig. 7F, S6H). There were also two clusters expressing markers more consistent with circulating populations of classical (CD14, S100A8, S100A9^38,39^ and non-classical monocytes (FCGR3A^38,39^). Looking at relative expression of *IL1RN* and *IL1R2* across different clusters and ages, we observed *IL1R2* to be preferentially expressed by mature macs, mono-Macs, cDC2 and LCs, without significant age-dependent trends (Fig. S6I). For *IL1RN*, however, preferential expression was noted in the mono-mac and mature macrophage clusters, with the highest expression for both cell types noted in the prenatal skin sample (Fig. 7G-H). Taken together, these observations suggest that monocyte-derived populations in human skin also are likewise key sources of IL-1 pathway repressors, especially with heightened potential for IL1-R2 and IL-1RA production during early skin development.

### Modulation of IL-1 signaling in early life restrains development of cutaneous type 17 responses

Based on the identification of IL-1RA producing monocytic cells in neonatal human and murine skin, we set out to test the model that depletion of neoMono leads to an increased type 17 immune response via unleashing of IL-1 bioavailability and subsequent signaling. To do so, we treated NeoΔMono mice with low dose recombinant IL-1Ra concurrently with anti-Gr1 administration, to compensate for the loss of IL-1 regulation by monocytes (Fig. 7I). Indeed, NeoΔMono mice that received IL-1Ra demonstrated significantly fewer IL-17A producing CD4^+^ Teff in the skin on day 21 (Fig. 7J). Notably, IL-1Ra treatment between day 4 and 14 was not sufficient to suppress IL-17A production by dermal γδ T cells on day 21 (Fig. S6J), and there was not a statistically significant effect on either CD4^+^ Teff and dermal γδ T cells in the SDLN (Fig. S6K). To further confirm that heightened IL-1R1 signaling in CD4^+^ T cells was contributing to their increased propensity for IL-17A production, we crossed *Cd4^cre^*with *Il1r1^fl/fl^* mice to achieve deletion of IL-1R1 on CD4^+^αβ T cells. We then subjected these mice and their *Il1r1^fl/fl^* littermates to our neonatal regimen of anti-Gr-1 treatment between day 4 and 14 (Fig. 7K).

*Cd4^cre^Il1r1^fl/fl^* mice demonstrated significantly lower percentages and numbers of Th17 cells in the skin on day 21 (Fig. 7L), whereas the effect was again less pronounced in the SDLN (Fig. S6L). Taken together these data suggest that neoMono contribute to negative regulation of IL-1 signaling in the developing tissue, thereby specifically and persistently constraining IL-17 production by skin CD4^+^ T cells.

## Discussion

Here we have identified a prominent postnatal wave of monocytes into skin as a key feature of normal early life development at this prominent body barrier site. Tissue accumulation of these monocytes early in life is critically facilitated by commensal microbes and requires Myd88-mediated signals. Whereas Ccr2^hi^Ly6C^hi^, so called “classical”, monocytes are typically recognized for their role in pro-inflammatory processes, we demonstrate a previously unappreciated regulatory role for them during healthy postnatal development. Depletion of neoMono in this window led to a sustained increase in IL-17A produced by CD4^+^ αβ T cells in the skin and draining lymph nodes, with a similar but more transient effect on dermal γδ T cells. Mechanistically the Th17 response was triggered by augmented upstream IL-1R1 signaling and sustained by the presence of commensal microbiota. Thus, neoMono accumulate in large numbers at the cutaneous interface and, via their enriched expression of IL-1 pathway repressors, help regulate type 17 responses and lower future susceptibility to skin conditions such as psoriasiform inflammation.

Our work demonstrating that commensal microbes facilitate early life skin seeding of CCR2^hi^Ly6C^hi^ monocytes adds to a growing literature regarding the unique immune composition in developing tissues and the role endogenous microbiota play in shaping this landscape^6,19,40^. Reduced neoMono in *Myd88^-/-^* but not *Il1r1^-/-^* skin further suggests that homeostatic TLR activation by commensals in early-life may provide key signals supporting their recruitment. While we were able to augment neoMono skin accumulation in germ-free mice by mono-colonization with *S. epi*dermidis, future studies are needed to disentangle the relative contribution of different bacteria and individual skin cell types in providing signals for skin neoMono recruitment, as well as the specific chemokines and receptors involved. Given the many skin cell types that express TLRs^41^ and the variety of signals that can contribute to monocyte recruitment^42–51^, it may be the case that there are multiple and redundant contributors.

Myeloid cells are key initiators of immune responses at barrier sites and as such contribute significantly to setting the immunological tone of these tissues. This has been best described for DC subsets and other antigen presenting cell types which tune tissue immune function in the skin and gut via their effects on T cells, both augmenting their production of key cytokines^22^ and also facilitating immune regulation via generation of Tregs^10,11,52^. In this study we report a previously unappreciated role for neoMono in skin immune regulation. In contrast to DCs, however, this immune regulation does not occur via augmented peripheral Treg generation but rather a direct effect on conventional CD4^+^ αβ T cells and dermal γδ T cells, namely limiting their production of the important barrier cytokine IL-17A. Depletion of neoMono had an impact on CD4^+^ αβ T cells which lasted several weeks before waning, as demonstrated by increased susceptibility of the NeoΔMono mice to psoriasiform inflammation at two but not six weeks post anti-Gr-1 treatment. While the increased IL-17A production was facilitated, at least in part, by the presence of skin microbes, it will be important for future studies to disentangle the antigenic specificities and targets of these dysregulated T cell responses and whether it is the overall burden of skin bacteria or the specific expansion of certain genera that is critical in driving the response.

IL-1 signaling is pivotal in mounting inflammatory responses to eliminate microbial infection and aid in physiological processes such as wound healing^53,54^. Suppression of this pathway by various cell types is critical for resolution of acute inflammatory responses^55,56^. The IL-1 receptor antagonist, or IL-1Ra, along with the decoy receptor IL-1R2 are two key molecules regulating IL-1R1 signaling. Loss of either can influence host susceptibility to infections ^57^, autoinflammatory disorders^58^ and other chronic inflammatory and autoimmune diseases^59,60^. The use of IL-1Ra (Anakinra) in the clinic to treat several skin and systemic disorders^60,61^ underscores the importance of the IL-1 pathway and its regulation during inflammation. Comparatively, little is known about its functional significance during homeostasis. Previous studies have detected intracellular IL-1Ra in healthy skin^62,63^. Our work highlights that neoMono and their mono-Mac progeny in neonatal skin are specifically enriched in IL-1Ra and IL-R2 expression as compared to other cell types or their adult skin counterparts. Our ability to find analogous mono-mac cells in human skin that demonstrate preferential early life expression of IL-1Ra suggest that this may be a more broadly conserved mechanism of immune regulation in developing tissue. The factors that elicit expression of these IL-1 pathway repressors in neoMono and why this is particularly heightened in the postnatal window remain key outstanding questions.

The attenuated Th17 response to neoMono depletion in the setting of IL-1Ra treatment or *Cd4^cre^Il1r1^fl/fl^* mice indicates that increased tissue availability of IL-1α and/or IL-1β has an effect via direct signaling on CD4^+^ T cells, as opposed to indirect effects via another intermediary population such as DCs. IL-1β has a well-documented role in promoting type 17 responses in T cells^53^, especially early in vivo Th17 differentiation via increased *Irf4* and *Rorc* expression^64^. IL-1R1 signaling on memory T cells, however, has also been shown to contribute to the heightened production of effector cytokines, including IL-17A, in previously differentiated cells^65^. The fact that we observed a polyclonal skin Th17 response that was heightened for several weeks but not indefinitely leaves open the question of whether altered differentiation and early life fate commitment of naïve CD4^+^ T cells is a central mechanism in our model. Further experiments designed to track subsets antigen-specific CD4^+^ T cells through this early life window through to adulthood would perhaps be helpful to further elucidate this possibility. In contrast, the effect of neoMono depletion on γδ T cell IL-17A production was quite transient and similar effects were not observed in ILCs based on our day 15 scRNA-seq analysis. While reversal effects in IL-17A production by γδ T cells was not anticipated in *Cd4^cre^Il1r1^fl/fl^*mice, we were surprised that their cytokine production in NeoΔMono was minimally impacted by IL-1Ra administration. This could be due to innate-like features of γδ T cells, with a lack of committed differentiation. However, we cannot exclude that some pathway other than IL-1R1 signaling is the main mechanism for IL-17A regulation in this cell type^66^. Separately, the lack of an effect for anti-Gr1 treatment in adult life, perhaps explained by the fact that monocytes are comparatively scarce in adult skin and express less IL-1Ra and IL-1R2, suggests that other cell populations might take over the role of IL-1 regulation during adult life.

That the elimination of neoMono leads to unleashing of IL-17A production by skin T cells has important implications for many physiological and pathological processes in skin. IL-17A production is critical in skin’s ability to fight acute bacterial and fungal infections. Other recent studies have demonstrated additional roles for IL-17 in critical tissue functions such as skin wound healing and nerve regeneration^67,68^. However, when unregulated, it is associated with key pathological processes in skin, namely psoriasis^69,70^, hidradenitis suppurativa^70,71^, some forms of atopic dermatitis^72^ among others^73^. The pleotropic effects of IL-17 suggest a need for tight regulation in the developing tissue to set the foundation for long-term skin health. Our work suggests that even in the absence of tissue inflammation, neonatal monocyte-mediated upstream regulation of IL-1 bioavailability serves as this rheostat for establishing a balanced type 17 lymphocyte compartment during a key developmental period.

The idea that monocytes can serve regulatory as well as proinflammatory roles is not unprecedented. In fact, this may be a key feature of “patrolling type” CX3CR1^hi^ CCR2^low^ monocytes in blood vessels. Additionally, Ly6C^hi^ monocytes have been shown to limit inflammation to gut commensal organisms after their recruitment to the site of intestinal infection via production of prostaglandin E2^74^. By comparison, our findings demonstrate a novel regulatory function for these Ly6C^hi^ neoMono in healthy tissue and one that is especially critical and heightened during postnatal life. This work emphasizes the importance of continuing to dissect fundamental mechanisms shaping tissue immunity in the early life window as these may uncover new opportunities to understand the persistent immune effects that can be imparted by early life exposures and their contribution to human health.

## STAR Methods

### RESSOURCES AVAILABILITY

#### Lead contacts

Further information and requests for resources and reagents should be directed to and will be fulfilled by the lead contacts, Miqdad Dhariwala Tiffany Scharschmidt

#### Material availability

All unique/stable reagents generated in this study are available from the Lead Contacts with a completed Materials Transfer Agreement.

#### Data and code availability

Schematics were made with Biorender (https://biorender.com). Rstudio v4 was used for scRNA-seq and CyTOF analysis. The scRNA-seq data generated in this study has been deposited into GEO and will be publicly available once this manuscript is accepted for publication.

### EXPERIMENTAL MODEL AND SUBJECT DETAILS

#### Experimental Animals

Wild-type C57BL/6 mice were originally purchased from Jackson Laboratories (Bar Harbor, ME), then bred and maintained in the UCSF specific pathogen-free (SPF) facility on the Parnassus campus for use in experiments. *Cd4^cre^* and *Il1r1^fl/fl^* mice were purchased from Jackson Laboratories and bred in-house. *Myd88^-/-^* and *Il1r1^-/-^* mice were obtained respectively from the labs of Averil Ma and Clifford Lowell at University of California, San Francisco (UCSF).

Gnotobiotic C57BL/6J mice (females and males, at indicated ages) were obtained from the UCSF Gnotobiotics core facility (gnotobiotics.ucsf.edu) and housed in gnotobiotic isolators for the duration of each experiment (Class Biologically Clean).

The age of all experimental animals is indicated in each figure. Littermates of the same sex were socially housed under a 12-hour light/dark cycle and randomly assigned to experimental groups whenever possible. Animal work was performed in accordance with the NIH Guide for the Care and Use of Laboratory Animals and the guidelines of the Laboratory Animal Resource Center at the University of California San Francisco, University Laboratory and Animal Resources at The Ohio State University along with Institutional Animal Care and Use Committee of both institutions.

#### Bacterial strain and culture conditions

*Staphylococcus epidermidis* (*S. epi*) strain Tü3298^75^ was used in this study and grown in tryptic soy broth at 37°C with shaking.

#### Methods Details

##### Heading Antibiotic treatments

For systemic antibiotic treatment in Figure 2, a cocktail of antibiotics including Ampicillin (1mg/ml), Metronidazole (1mg/ml). Sulfamethoxazole (0.8mg/ml) and Trimethoprim (0.16mg/ml) was dissolved in the drinking water and supplied to pregnant dams. To reduce the microbial burden in the environment, cages were changed every other day for the antibiotic treated mice whereas controls cages were changed once a week as per conventional housing and husbandry policies. Control animals underwent mock cage changes. For topical antibiotic treatments in Figure 4, commercially available Neosporin was applied on the ear skin of mice with cotton swab applicators at the indicated ages and intervals. Control mice were treated with Vaseline.

##### Antibody injections

Mice were injected with either the anit-Gr-1 (100ug) (Bioxcell, InvivoMAb Catalog No. BE0075), Ly6G (100ug) (Bioxcell, InvivoMAb Catalog No. BE0075-1), Ccr2 (2mg/kg, provided by Matthias Mack and Amar Nijagal) or isotype control (100ug) (Bioxcell, InvivoMAb Catalog No. BE0090) via the intraperitoneal route at the indicated time points.

##### Mouse skin processing

Back skin was shaved and harvested, lightly defatted, and then minced with scissors to a fine consistency before tissue digestion in 3 mL complete RPMI (RPMI plus 10% fetal calf serum, 1% penicillin-streptomycin, β-mercaptoethanol, glutamate, sodium pyruvate, HEPES and non-essential amino acids) then supplemented with 2 mg/mL collagenase XI, 0.5 mg/mL hyaluronidase, 0.1 mg/mL DNase I (digestion media). Ear pinna skin was split and dorsal and ventral sheets were placed in PBS. Ear skin was then processed similarly to the back skin in a 1ml total volume of digestion media. Digested skin samples were then incubated, with shaking, at 37°C for 45 minutes before quenching with 10 mL of complete RPMI media and shaking by hand for 30 seconds. Skin cell suspensions were filtered through sterile cell strainers (100 μm followed by 40 μm).

##### Mouse lymph node processing

Skin draining lymph nodes were harvested and then processed by gentle grinding with a sterile plunger over sterile wire mesh in 1 mL of complete RPMI media to generate a single cell suspension for further analyses.

##### Cell counting

All tissues were re-suspended in 1 mL PBS and 25 μL of cell suspension was mixed with 25 μL of AccuCheck counting beads (Invitrogen, Catalog No. PCB100) for calculating absolute numbers of cells.

### Flow Cytometry

#### Antibody Staining of Myeloid Populations

Cells were stained in PBS with a Live/Dead marker (Ghost Dye Violet510, Tonbo Biosciences, Catalog No. 13-0870-T100), followed by Fc block (Fisher Scientific, Catalog No. BDB564219) for 15 minutes each at 4°C. Surface antibodies were added in blocking solution and cells stained for 60 minutes at 4°C. Cells were then washed and fixed in Paraformaldehyde (PFA) for 30 minutes at 4°C.

#### Antibody Staining of Lymphoid Populations

Cells were stained in blocking solution with a Live/Dead marker (Ghost Dye Violet510, Tonbo Biosciences) for 30 minutes at 4°C. For intracellular staining, cells were fixed and permeabilized using the Foxp3 staining kit (eBioscience, Catalog No. 00-5523-00) buffer for 30 minutes at 4°C then stained in permeabilization buffer for 30 minutes at 4°C.

#### Cell Re-stimulation for Cytokines

After isolation, skin or SDLN single cell suspensions were stimulated with Tonbo kit (1/100 in 1 mL) for 3-6 hours before being processed for staining. A subset of cells was incubated in Brefeldin A to be used as an unstimulated control for gating.

#### Flow cytometry and analyses

Stained cells were run on a Fortessa (BD Biosciences) in the UCSF Flow Cytometry Core. Flow cytometry data was analyzed using FlowJo software (FlowJo, LLC).

#### Mass cytometry and analyses

Single cell suspensions were incubated for 1 minute with 25 mM Cisplatin (Sigma-Aldrich, P4394) to allow subsequent cell viability measurement, then fixed in 1.5% paraformaldehyde (Electron Microscopy Sciences) and frozen down in the presence of 10% di-methyl sulfoxide (DMSO) and 10% bovine serum albumin (BSA) at stored at 80°C for subsequent staining. Individual vials were thawed and 2×10^6^ cells were barcoded using the Cell-ID 20-Plex Pd Barcoding Kit (Fluidigm, product number 201060) for 15 minutes at room temperature. Palladium based barcoding was used as this has been reported to not only enable simultaneous staining of up to 20 different samples but also enable doublet discrimination^76^. All the barcoded samples were then combined into a single tube and cells were stained with metal conjugated antibodies, purchased either from BD Biosciences or Fluidigm. Surface antigens were stained in cell staining media (0.5% BSA, 0.02% Sodium azide in PBS) for 30 minutes at room temperature with gentle agitation. For intracellular staining, cells were permeabilized using the FOXP3-staining buffer kit (eBioscience) and then incubated for another 30 minutes at room temperature with intracellular antibodies in the presence of the FOXP3-staining kit permeabilization buffer. Cells were stored overnight at 4°C in a buffer containing iridium and then run on a Helios CyTOF system (Fluidigm) in the UCSF, Parnassus flow cytometry core facility.

CyTOF data analysis was performed using the *Cyclone* R package^77^. To identify cell clusters, unsupervised clustering was implemented using the *FlowSOM* algorithm with an optimized 6 by 5 grid to generate 30 clusters^78^. The resulting cell clusters were manually annotated with cell type based on the lineage marker expression profile of each cluster. The markers B220 and NK1.1 demonstrated poor staining quality based upon downstream analysis and were consequently excluded from the unsupervised clustering algorithm. Uniform Manifold Approximation and Projection (UMAP) dimensionality reduction was performed using the *uwot* R package and used to visualize the relationship between cell clusters^79^. To broadly gauge the effect of age, principal-component analysis (PCA) was performed separately on cluster lineage marker expression as well as the percentage of cells per cluster using the prcomp function from the *stats* R package^80^.

#### 16S rRNA sampling

Sterile foam tip applicators were dipped in sterile Yeast lysis buffer (*Epicentre Biotechnologies; product number MPY80200*) and rubbed on the back skin of Gr-1 treated or control littermate mice at postnatal day 15. Tips of these applicators were then stored in Eppendorf tubes in 100ul of yeast lysis buffer. Tubes were stored at -80°C and then shipped to SeqCenter for DNA extraction and 16s rRNA sequencing.

#### DNA extraction

DNA was extracted at the SeqCenter facilities using the ZymoBIOMICS^TM^ DNA Miniprep Kit. Briefly, lysis buffer was transferred into the ZR BashingBead™ Lysis Tubes and mechanically lysed using the MP FastPrep-24™ lysis system with 1 minute of lysis at maximum speed and 5 minutes of rest for 5 cycles. Samples were then centrifuged at 10,000rcf for 1 minute. 400μl of supernatant was transferred from the ZR BashingBead™ Lysis Tube to a Zymo-Spin™ III-F Filter and centrifuged at 8,000rcf for 1 minute. 1,200 μl of ZymoBIOMICS™ DNA Binding Buffer was added to the effluent and mixed via pipetting. 800μl of this solution was transferred to a Zymo-Spin™ IICR Column and centrifuged at 10,000rcf for 1 minute. This step was repeated until all material was loaded onto the Zymo-Spin™ IICR Column. DNA bound to the Zymo-Spin™ IICR Column was washed 3 times with 400μl and 700μl of ZymoBIOMICS™ DNA Wash Buffer 1 and then 200 μl of ZymoBIOMICS™ DNA Wash Buffer 2 with a 1-minute spin down at 10,000rcf for each, respectively. Washed DNA was eluted using 75μl of ZymoBIOMICS™ DNase/RNase Free Water following a 5-minute incubation at room temperature and a 1-minute spin down at 10,000rcf. The Zymo-Spin™ III-HRC Filter was prepared using 600 μl of the ZymoBIOMICS™ HRC Prep Solution and a centrifugation at 8,000rcf for 3 minutes. Eluted DNA was then purified by running the effluent through the prepared Zymo-Spin™ III-HRC Filter. Final DNA concentrations were determined via Qubit.

#### 16S sequencing

Samples were prepared using Zymo Research’s Quick-16S kit with phased primers targeting the V3/V4 regions of the 16S gene. The specific primer sequences are listed in the STAR methods section. Following clean up and normalization, samples were sequenced on a P1 600cyc NextSeq2000 Flowcell to generate 2×301bp paired end (PE) reads. Quality control and adapter trimming was performed with bcl-convert (v4.2.4).

#### 16S Data analyses

For the sequence analysis DADA2 (https://benjjneb.github.io/dada2/bigdata_paired.html) was run on reads following a modified big data workflow, using a one-by-one pseudo-pooling approach for ASV inference and chimera detection, with the following parameters (anything not listed was left at defaults):

| **filter and trim** | **forward** | **reverse** |
| --- | --- | --- |
| **parameter** | **reads** | **reads** |
| maxN | 0 | 0 |
| maxEE | 2 | 3 |
| trimleft | 17 | 21 |
| truncLen | 270 | 250 |

Taxonomy was determined using SILVA’s non-redundant at 99%, with species, database constructed for DADA2. ASVs and their reads removed from downstream analyses due to either lacking Phylum-level taxonomic inference, affiliation with plastids (chloroplast, mitochondria), extremely low abundances, or identification as a likely contaminant using the decontam R package (https://benjjneb.github.io/decontam/vignettes/decontam_intro.html). Differential abundance testing using ANCOM-BC (https://www.bioconductor.org/packages/release/bioc/vignettes/ANCOMBC/inst/doc/ANCOMBC.html) at each taxonomic level. For all taxonomic-level analyses, the read counts/relative abundances and number of ASVs are aggregated without the unassigned members. The NMDS was solved from Bray-Curtiss dissimilarity matrices, the statistical support for similarity between the defined groups as estimated by ANOSIM and ADONIS2 (i.e., PERMANOVA), and significant (R > 0.1, p < 0.05) fits to the ordination of genera were determined using envfit, all implemented in Vegan (https://cran.r-project.org/web/packages/vegan/index.html). For ASVS with mean relative abundance <1*10^-5 were excluded from these analyses, though they were not identified as likely contaminants with low abundance may introduce excessive noise. For the ranked-abundance curves, no ASVS were filtered out due to low abundances.

#### Single cell RNA sequencing

Live singlet immune cell populations (CD45^+^) from the back skin of postnatal D15 mice treated with anti-Gr-1 antibody or isotype controls were sorted using a BD FACSAria II with a 100 μm nozzle into RPMI medium supplemented with 10% FBS. Cells were sorted from pooled tissue from 4-7 mice and different conditions were run on separate lanes of a 10X Chromium chip with 3′ v.2 chemistry (10X Genomics) as per the manufacturer’s instructions by the UCSF Institute for Human Genetics Sequencing Core. Libraries were sequenced on an Illumina NovaSeq 6000. FASTQ files were aligned to the mm10 reference genome, and barcode matrices were generated using CellRanger 2.1.

Downstream data analysis, including clustering, visualizations and exploratory analyses, were performed using Seurat 4.0. Cells with <200 features or more than 5000, or >10% mitochondrial genes, and >30% ribosomal genes were filtered out during preprocessing. Principal component analysis (PCA) and uniform manifold approximation and projection (UMAP) were run on the RNA data, and clustering was generated using the first 25 principal components and a resolution of 1. Fast gene set enrichment analyses (fGSEA) for the IL-1 pathway was performed using the genes *Adamts4, Adamts5, App, Casp8, Col3a1, Csf2, Cxcl2, Ifng, Il17a, Il1a, Il1b, Il22, Il6, Mapk8, Mmp13, Mmp14, Mmp3, Mmp8, Mmp9, Nos2, S1008, Sele, Smad6, Tnf, Il1rn, Il1r2*.

<u>Human</u> raw sequencing data were aligned and quantified against the human reference genome GRCh38 using CellRanger (10X Genomics). Cells with more than 10-20% mitochondrial and 60% ribosomal content were excluded. Donor samples were demultiplexed based on sequenced SNPs using the Demuxlet^81^ software. Intrasample doublets were removed using DoubletFinder^82^ . Counting matrix was loaded into Seurat, where differentially expressed genes (DEGs) were identified using the FindMarkers function. Cell clusters were manually annotated and grouped into major immune cell types, including B-cells, T-cells, NK-cells and myeloid cells, and quantified for each donor. The mean and standard deviation of cell abundances were calculated for the two adult donors. Myeloid cell clusters were subset, normalized, scaled, and PCAs, neighbor identification, clustering, and UMAP were performed. DEGs and cluster annotations were performed as described above.

#### Quantitative RT-PCR

Indicated populations were sorted into RLT Plus lysis buffer (Qiagen) and stored at -80°C, then processed using Allprep DNA/RNA micro kit (Qiagen) per manufacturer’s protocol. For qPCR analyses, RNA was reverse transcribed using SuperScript III cDNA synthesis kit (ThermoFisher) and amplified using Power SYBR Green PCR master mix (ThermoFisher). *Il1rn* primers: 5′-TAGACATGGTGCCTATTGACCT -3′, forward, and 5′-TCGTGACTATAAGGGGCTCTTC -3′, reverse. *Il1r2* primers: 5′-GATCCAGTCACAAGGGAGGA -3′, forward, and 5′-CCAGGAGAACGTGGAAGAGA -3′, reverse. *Rsp17* gene was used as a <u>housekeeping gene</u> and amplified with the following primers: 5’-ATTGAGGTGGATCCCGACAC -3’, forward; 5’-TGCCAACTGTAGGCTGAGTG -3’, reverse.

#### Quantification and statistical analyses

The number of mice per group and independent experimental repeats are annotated in each of the corresponding figure legends. Mice were allocated randomly into experimental groups after matching for age and littermates were used as controls as best as possible, wherever applicable. Statistical significance was calculated using the GraphPad (Prism) 9.3.1 software and specifics for each experiment are indicated in the corresponding figure legends. Data presented here always demonstrate the statistical mean.

### Human skin samples processing

#### Human donor collection

A human prenatal donor (22-23 gestational weeks) was obtained from pregnancy terminations conducted with maternal written informed consent, in accordance with the approval from and under the guidelines of the UCSF Research Protection program. Samples were excluded if there was evidence of maternal infection, intrauterine demise, or known/suspected chromosomal abnormalities. Pediatric (16 years old) and adult (33 and 36 years old) donors were obtained from consenting organ donors under the UCSF VITAL Core protocol (IRB No 20-31618), in collaboration with Donor Network West. All donors were confirmed negative for HIV, HBV and HCV. Skin samples were collected by UCSF transplant surgeons and preserved in UW solution on ice until processing (<24 hours).

#### Preparation of skin single cell suspension

Samples were processed using a dermatome to isolate the top layer of the epidermis at a thickness of 0.6 µM and subsequently cut into pieces no larger than 4mm by 4mm. Skin pieces were cryopreserved with FCS with 10% DMSO at - 80°C until thawing and processing. <u>Tissue pieces were thawed with complete media (RPMI 1640, 10% FCS, 1% PSG, 1% HEPES) and rinsed with fresh complete media to thoroughly remove residual DMSO from the freezing media.</u> Skin tissue was digested in resting media (89% RPMI 1640, 10% FCS and 1% Pen-Strep-Glut) containing 750 ug/ml Collagenase IV (Thermo Fisher) and 20 ug/ml DNase I (Millipore Sigma) for 2 hours at 37°C. Cells were filtered through a 100 µm strainer (Corning), centrifuged at 500g for 5 minutes at 4°C, counted, and resuspended for flow cytometry staining.

#### Flow cytometry

Cells were transferred to a 96-well v-bottom plate and stained with LIVE/DEAD Fixable Aqua (Invitrogen, 1:500 in 1X PBS, 20 million cells per 1mL). Cells were washed with 150uL of sort buffer (2mM EDTA, 2% FBS in 1X PBS), centrifuged at 500g for 5 minutes at 4°C, and incubated with the antibody surface cocktail (STAR Table 1) for 30 minutes at 4°C. Cells were washed with 150uL of sort buffer and centrifuged at 500g for 5 minutes at 4°C. Live CD45+ cells were sorted with FACSAria Fusion Flow Cytometer.

#### scRNA-seq sample processing

Cells from four donors were evenly pooled (approximately 14K cells per donor) and encapsulated using the Chromium Next GEM Single Cell 5’ Reagent Kits V2 (Dual Index) according to the manufacturer’s instructions (10X Genomics). Libraries were amplified with 13 cycles of cDNA amplification. Sequencing was performed on the NovaSeq X Plus instrument at the UCSF Center for Advanced Technology. To identify SNPs for each donor, total RNA was extracted from the spleen or mesenteric lymph nodes samples using the RNeasy Micro Kit (QIAGEN), following the manufacturer’s instructions. RNA sequencing was performed with 150 bp paired-end reads at 30 million reads per sample on the NovaSeq instrument at the UCSF Genomics Facility.

## Acknowledgements

We would like to thank Drs. Michael Rosenblum, Ari Molofsky, Clifford Lowell and Abul Abbas for helpful discussions. We thank Drs. Matthias Mack and Amar Nijagal for providing the anti-Ccr2 antibody. We thank Dr. Clifford Lowell and Dr. Clare Abram for providing the *Il1r1^-/-^*mice and Dr. Averil Ma for providing the *Myd88^-/-^* mice. We thank Yongmei Hu for mouse husbandry along with Dolores Espiritu and Minerva Espiritu for lab infrastructure. We thank Robin Schwartz for her help with mouse genotyping and lab infrastructure at OSU. We thank and acknowledge members of the HSW12 community for helpful discussions. We thank Jessie Turnbaugh and the UCSF Gnotobiotics core for experimental help with Germ-free mice. We acknowledge UCSF’s PFCC (RRID:SCR_018206) for assistance in flow cytometry and cell sorting, supported by NIH grants P30 DK063720 and 1S10OD021822-01. We acknowledge the UCSF Genomics CoLab in supporting generation of scRNA-seq data. M.O.D. is supported by NIAMS K99AR079554. This work was funded by NIAID DP2AI144968, NIAMS R01AR080034 and an ImmunoX CoLabs Co-Project.

## Author Contributions

M.O.D. and T.C.S. drafted the manuscript and figures. M.O.D. performed the experiments and analyzed the data. A.M.D., H.R.V., A.W., I.H., S.V., and K.J.H. assisted in bioinformatic data analyses. R.O.R., A.M.D., J.N.O., C.E., A.W., V.M.T., R.B.C., O.T.O., A.E.T., J.T., A.K.A., J.M.L., and G.R.M. assisted in performing experiments. J.H., J.G., R.R., G.K.F, M.H.S. and A.J.C provided resources and input on experimental design and data interpretation. T.C.S. and M.O.D. oversaw all study design and data analyses. All authors discussed results and commented on the manuscript.

## Inclusion and diversity

Multiple authors of this paper self-identify as an underrepresented ethnic minority in their field of research or within their geographical location. Multiple authors of this paper self-identify as a gender minority in their field of research. One or more of the authors of this paper received support from a program designed to increase minority representation in their field of research. While citing references scientifically relevant for this work, we also actively worked to promote gender balance in our reference list.

