## Supplemental Materials for "Commensal myeloid crosstalk in neonatal skin regulates cutaneous type 17 inflammation"

### Supplementary Materials

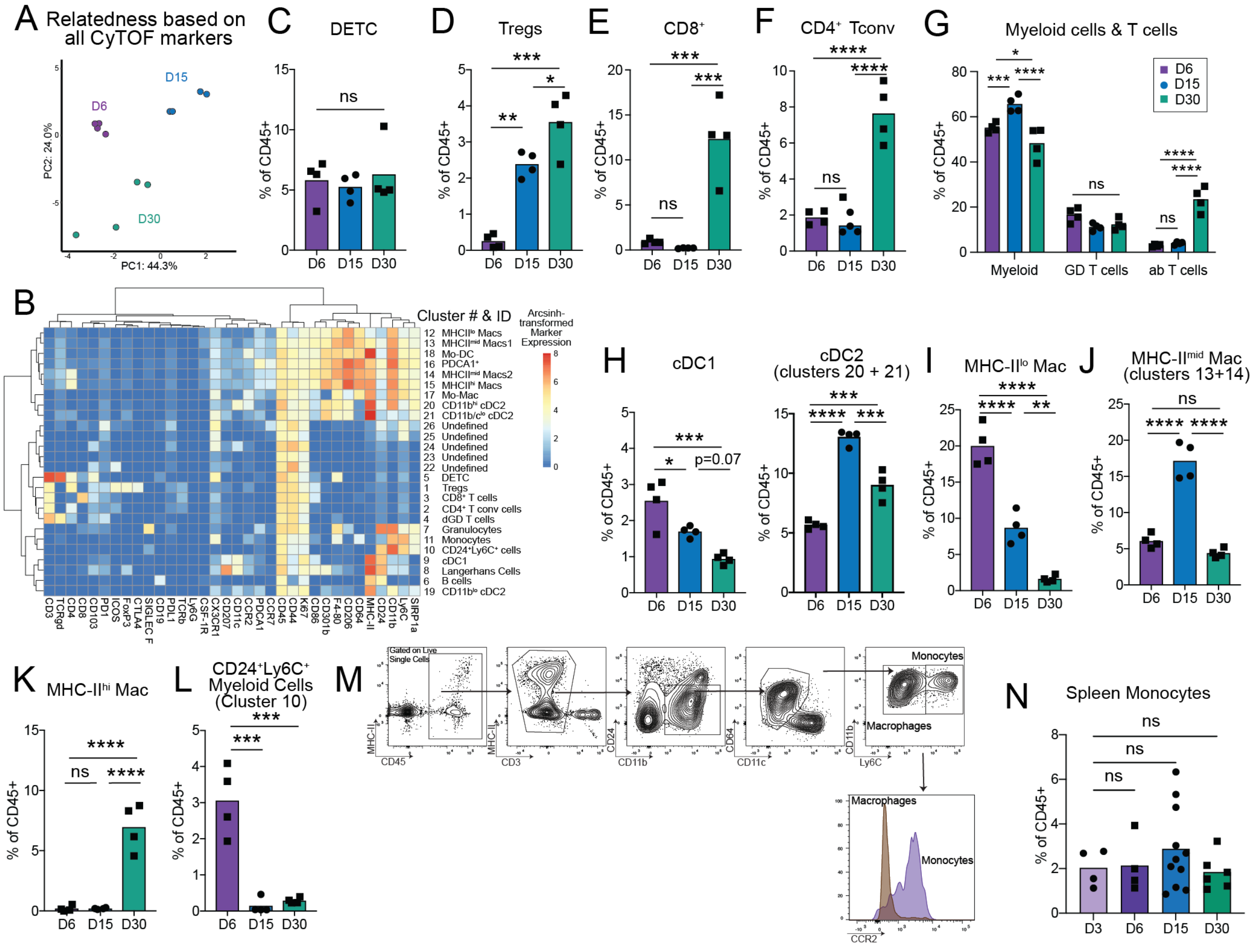


**Figure S1: Related to main Figure 1** (A) Principal component analysis (PCA) plot demonstrating the distribution of the cutaneous CD45$^{+}$ cellular compartment based on marker relatedness. Each dot represents a single mouse at the indicated age. (B) Heat map demonstrating CyTOF marker expression profiles across all immune clusters in postnatal Day 6, Day 15 and Day 30 skin. (C-H) Quantification of (C) Dendritic Epidermal T cells (DETC), (D) Tregs, (E) CD8$^{+}$ T cells, (F) CD4$^{+}$ T conventional cells, (G) pooled myeloid, $\gamma\delta$ T cell and $\alpha\beta$ T cells clusters, (H) Dendritic Cells, and (I-L), Macrophages as measured by CyTOF analyses of Day 6, Day 15 and Day 30 skin. (M) Gating strategy used for flow cytometric evaluation of monocytes in murine skin with a histogram demonstrating CCR2 expression levels on cutaneous monocytes and macrophages. (N) Flow cytometric quantification of monocytes in the spleen of postnatal Day 3, 6, 15 and 30 old mice. For CyTOF assays an n=4 mice were used at each time point, for flow cytometry assays an n>4 mice were used at each time point. Statistical significance was determined by a One-Way ANOVA or Two-Way ANOVA (G) where * is $p<0.05$ ** is $p<0.01$ *** is $p<0.001$ and **** is $p<0.0001$.

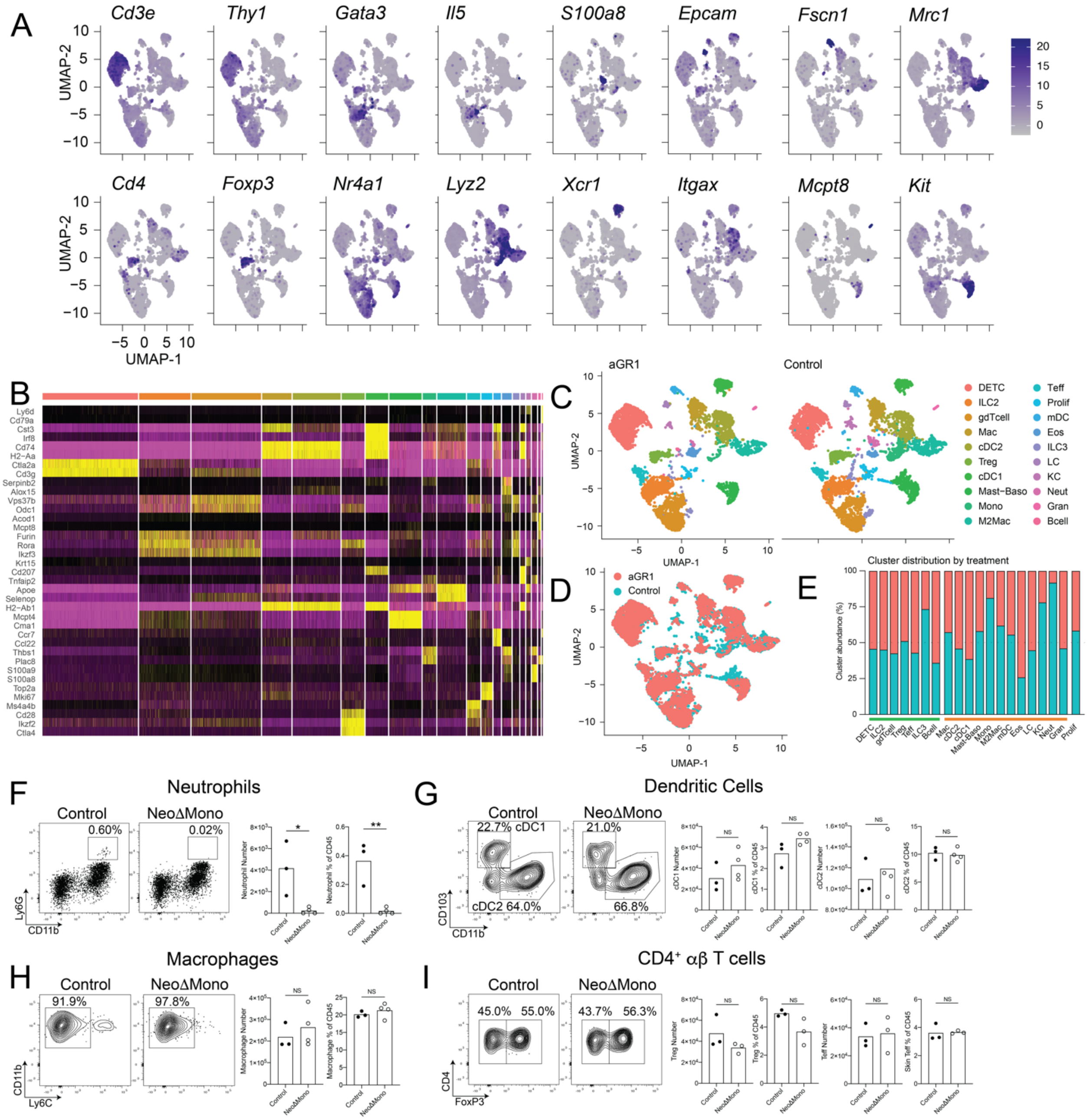


**Figure S2: Related to main Figure 3** (A) UMAPs demonstrating expression profiles of indicated genes in skin immune clusters of NeoΔmono and control mice. (B) Heatmap of genes used to define clusters in Fig. 3D. (C-D) Individual and overlaid UMAPs demonstrating immune cell clusters in the skin of anti-Gr-1 treated NeoΔmono and isotype treated control mice. (E) scRNA sequencing based quantification of relative cluster abundance in skin of anti-Gr-1 treated NeoΔmono and isotype treated control mice. (F-I) Representative flow cytometry plots and quantification of neutrophils (F), dendritic Cells (G), macrophages (H) and CD4⁺ T cells (I) in the skin of NeoΔmono and control mice on postnatal Day 15 of life. All scRNA-seq analyses were performed post normalizing clusters across both treatment groups. In F-I, statistical significance was determined by a two-tailed Student's t test where * is $p<0.05$ and ** is $p<0.01$.

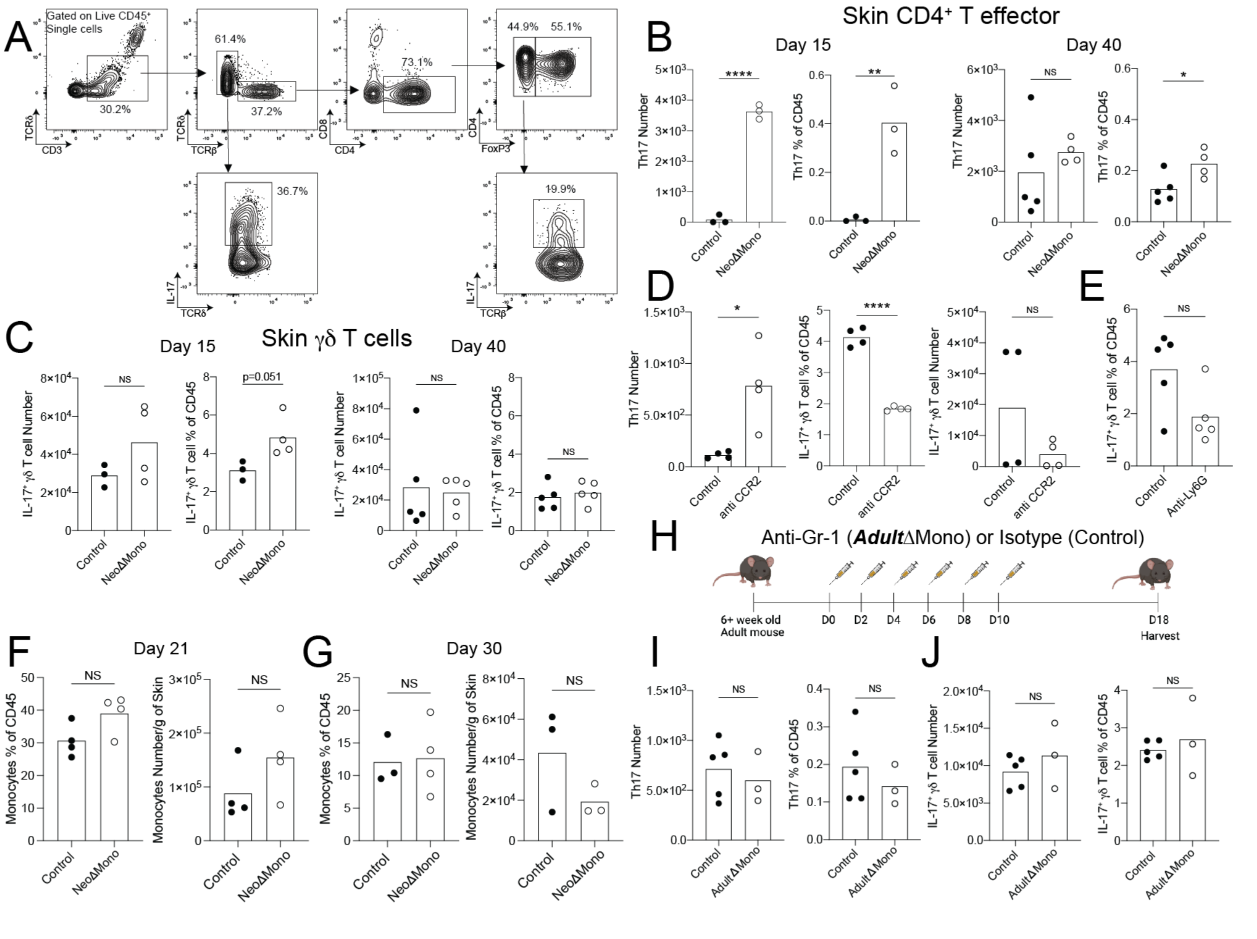


**Figure S3: Related to main figure 4** (A) Flow cytometric gating strategy to examine IL-17 producing γδ T cells and CD4⁺ αβ T effector cells in the skin. (B-C) Quantification of IL-17 producing CD4⁺ αβ T effector cells (B) and γδ T cells (C) in the skin of NeoΔmono and control mice at postnatal Day 15 and Day 40. (D-E) Quantification of numbers of CD4⁺ T effector cells producing IL-17 in the skin of mice treated with anti-CCR2 antibody (D) or anti-Ly6G antibody (E). (F-G) Quantification of numbers of monocytes present in the skin of NeoΔmono and control mice at postnatal day 21 and day 30. (H) Monocytes were depleted in >8week old adult mice by treatment with the anti-Gr-1 antibody (*Adult*Δmono mice) and controls were treated with isotype antibody. Skin from these mice were then processed for flow cytometry at indicated time points. (I-J) Quantification of IL-17A producing CD4⁺ αβ T effector cells (I) and γδ T cells (J) in the skin of *Adult*Δmono and control mice. Data here are one representative of at least two independent experiments. Statistical significance was determined by a two-tailed Student's t test where * is p<0.05, ** is p<0.01 and **** is p<0.0001

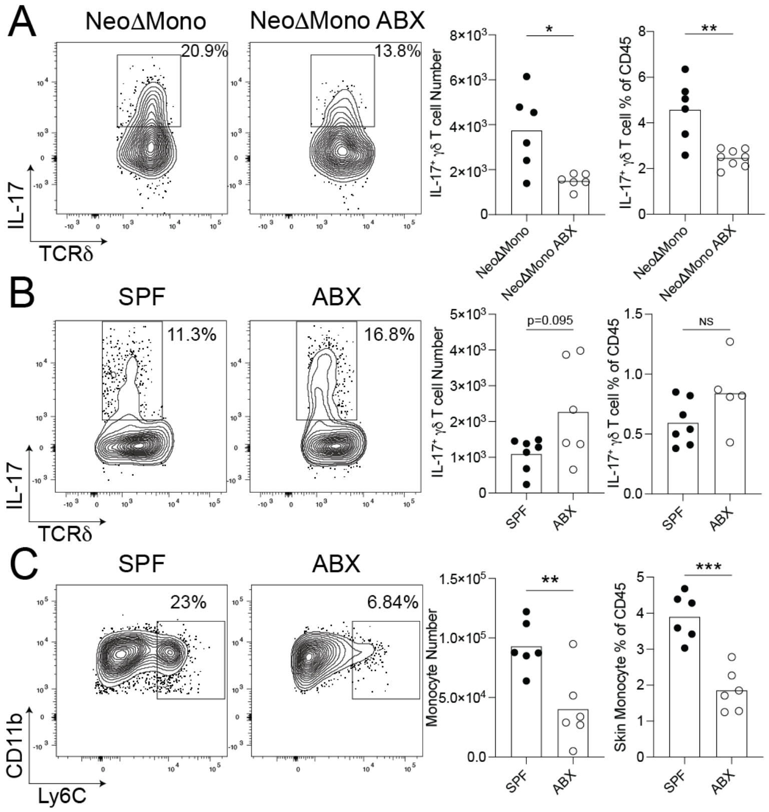


**Figure S4: Related to main figure 5 (A)** Representative flow cytometry plots and quantification of numbers of IL-17A-producing dermal γδ T cells in the ear skin at day 21 from NeoΔMono mice treated with topical Neosporin and frequent cage changes or controls treated with vehicle and subjected to mock cage changes. (B-C) Representative flow cytometry plots and quantification of numbers of IL-17-producing dermal γδ T cells (B) and (C) monocytes in the back skin of wild-type mice treated with systemic antibiotics and frequent cage changes from birth or untreated age-matched controls subjected to mock cage changes. Data in A are pooled from two of three representative experiments, data in B-C is one representative of 2 independent experiments. Statistical significance was determined by a two-tailed Student's t test where * is $p<0.05$, ** is $p<0.01$, and *** is p<0.0)1.

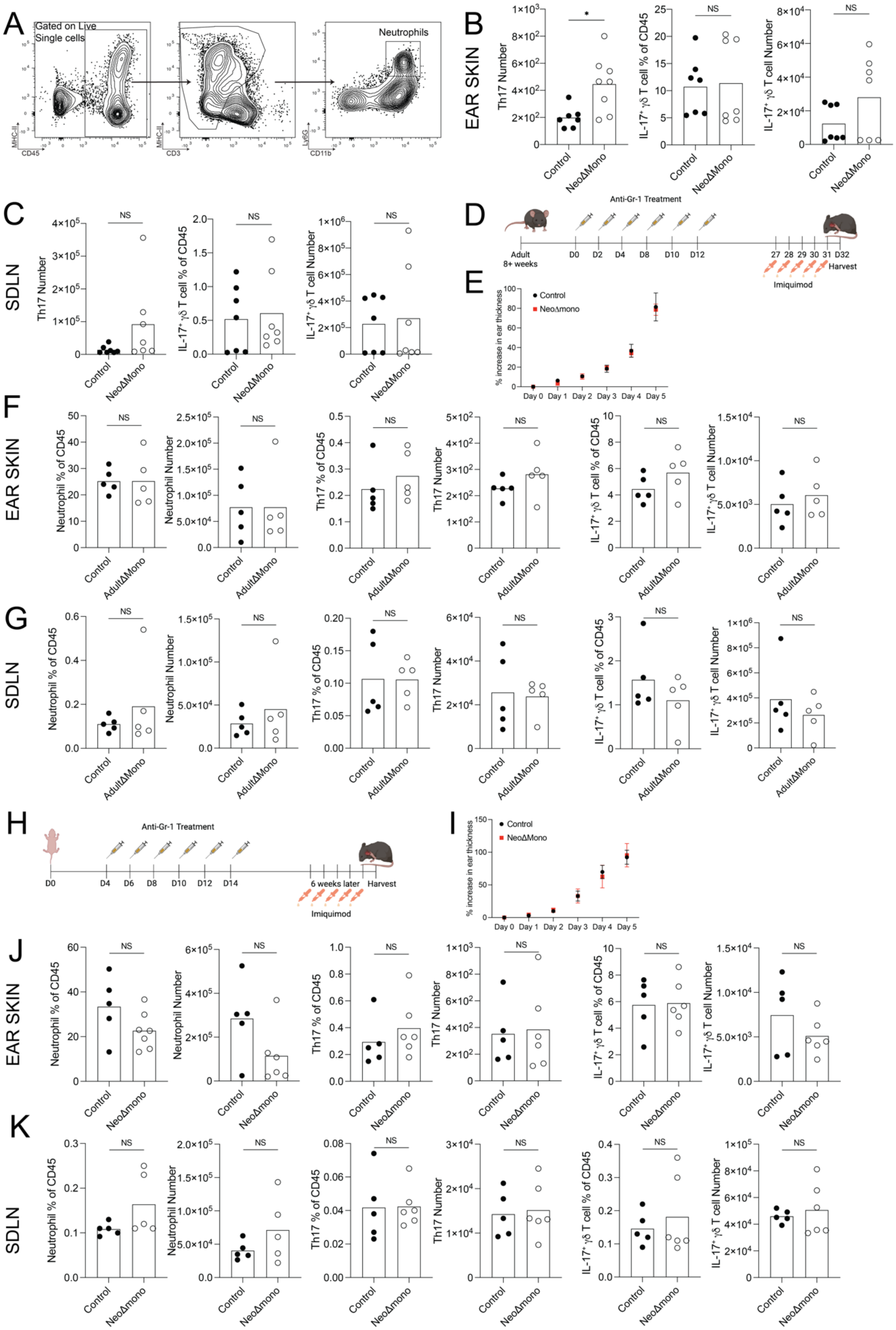

A
Gated on Live Single cells
Neutrophils
B
EAR SKIN
Th17 Number
IL-17⁺ γδ T cell % of CD45
IL-17⁺ γδ T cell Number
Control
NeoΔMono
NS
C
SDLN
D
Anti-Gr-1 Treatment
Adult 8+ weeks
D0 D2 D4 D8 D10 D12
27 28 29 30 31 D32
Harvest
Imiquimod
E
% increase in ear thickness
Control
NeoΔmono
Day 0 Day 1 Day 2 Day 3 Day 4 Day 5
F
Neutrophil % of CD45
Neutrophil Number
Th17 % of CD45
AdultΔMono
G
H
D0 D4 D6 D8 D10 D12 D14
6 weeks later
I
J
K

**Figure S5: Related to main Figure 6**

(A) Flow cytometric gating strategy to examine neutrophils in the skin and draining lymph nodes. (B-C) Quantification of IL-17A producing $CD4^+$ Teff cells and $\gamma\delta$ T cells in the ear skin (B) and draining lymph nodes (SDLN) (C) of Neo∆mono and control mice treated with 5% imiquimod cream. (D-G) Monocytes were depleted in adult mice with the anti-Gr-1 antibody (Adult∆Mono mice) and controls were treated with isotype antibody. The ears of these mice were treated daily for five days with 5% Imiquimod cream then one day later ear skin and lymph nodes were harvested. (E) Quantification of relative increase in ear thickness as measured with digital calipers during imiquimod treatment in Adult∆Mono versus control mice. (F-G) Quantification of neutrophils, $CD4^+$ Th17 cells, and IL-$17^+$ $\gamma\delta$ T cells in the ear skin (F) and ear draining lymph nodes (G). (H-K) Monocytes were depleted in neonatal mice with the anti-Gr-1 antibody (Neo∆Mono mice) and controls were treated with isotype antibody. The ears of these mice were treated daily for five days with 5% Imiquimod cream, 6 weeks after the final Gr-1 treatment, then one day later ear skin and lymph nodes were harvested. (I) Quantification of relative increase in ear thickness as measured with digital calipers during imiquimod treatment in Neo∆Mono versus control mice. (J-K) Quantification of neutrophils, $CD4^+$ Th17 cells, and IL-$17^+$ $\gamma\delta$ T cells in the ear skin (J) and ear draining lymph nodes (K). Data in B-C is pooled data from two independent experiments. Data in E-K is from one of two independent experiments. Statistical significance was determined using a two-tailed Student's t test where * is $p<0.05$.

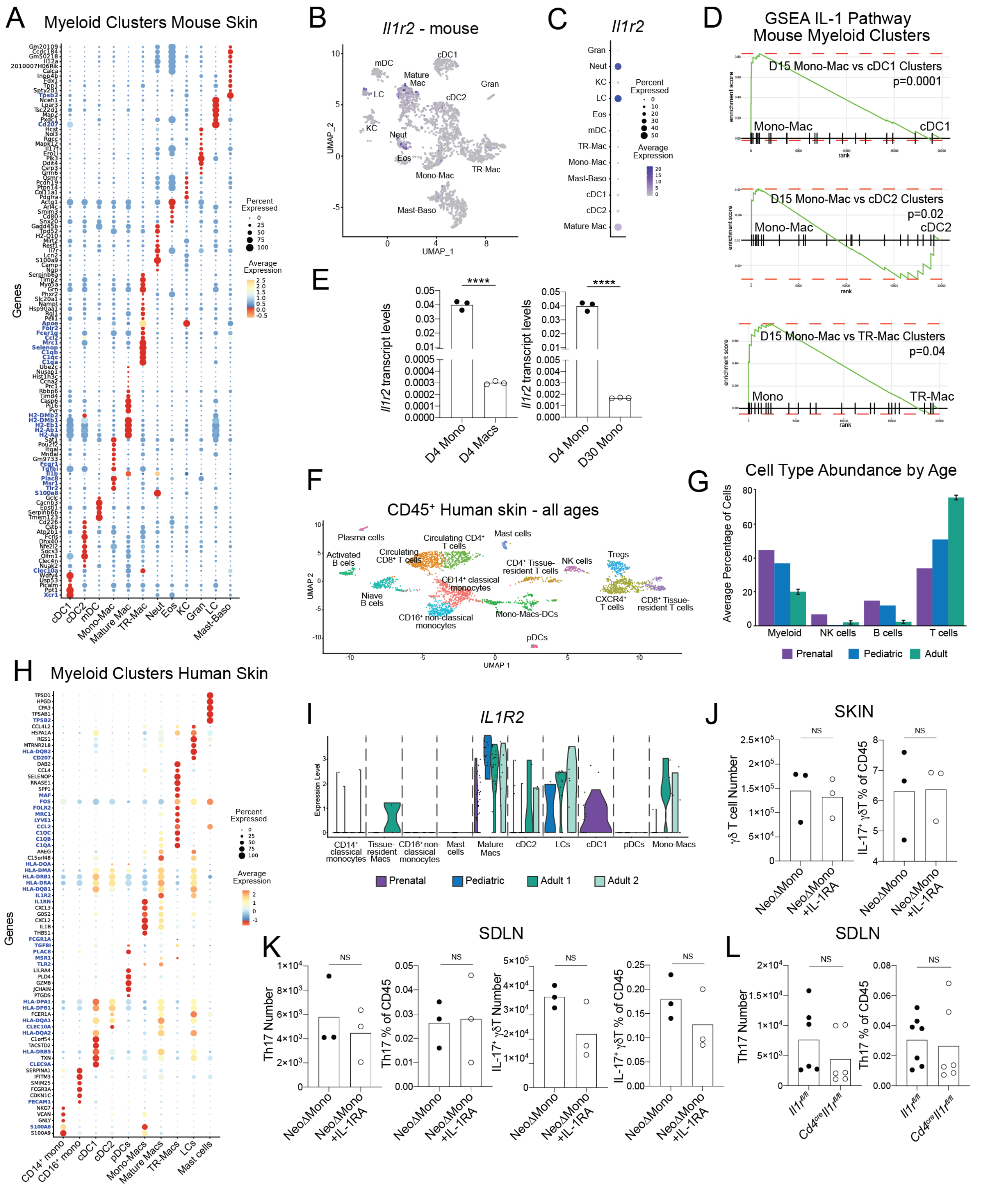


**Figure S6: Related to main Figure 7** (A) Key differentially expressed genes by cluster from scRNAseq of myeloid cells from D15 mouse skin. (B) Intensity plot showing *Il1r2* expression across murine skin myeloid clusters. (C) Quantification of *Il1r3* expression by normalized average expression and percentage of expressing cells for individual myeloid clusters. (D) Gene Set Enrichment Analyses (GSEA) for IL-1 pathway genes comparing postnatal Day 15 skin mono-mac and cDC1, mature mac and cDC2 clusters.

(E) qPCR analyses for *Il1r2* transcript levels comparing monocytes versus macrophages sorted from postnatal Day 4 skin and monocytes sorted from postnatal Day 4 versus Day 30 skin. (F) UMAP of all clusters from scRNA-seq analysis of $CD45^+$ cells sorted from human skin – combining cells from all four donors. (G) Enumeration of relative abundance of key cell types as a percentage of $CD45^+$ cells by age from scRNAseq data. (H) Key differentially expressed genes used for human myeloid cluster annotation. (I) Violin plot showing *IL1R2* expression across human skin myeloid clusters by age. (J) Quantification of numbers of IL-17A producing $\gamma\delta$ T cells in the skin (K) and SDLN (K) along with $CD4^+$ $\alpha\beta$ T effector cells in the SDLN (K) of Neo$\Delta$mono mice treated with recombinant IL-1Ra or vehicle control. (L) Quantification of numbers of $CD4^+$ T effector cells producing IL-17A in the SDLN of $Cd4^{cre}Il1r1^{fl/fl}$ or control $Il1r1^{fl/fl}$ mice treated with anti-Gr-1 antibody in the neonatal window. Statistical significance was determined using a two-tailed Student's t test where **** is p<0.0001. Data in E-F are from one of three independent experiments and data in L is pooled from two of three independent experiments.
